# Activation of Polycystin-1 Signaling by Binding of Stalk-derived Peptide Agonists

**DOI:** 10.1101/2024.01.06.574465

**Authors:** Shristi Pawnikar, Brenda S. Magenheimer, Keya Joshi, Ericka Nevarez Munoz, Allan Haldane, Robin L. Maser, Yinglong Miao

## Abstract

Polycystin-1 (PC1) is the membrane protein product of the PKD1 gene whose mutation is responsible for 85% of the cases of autosomal dominant polycystic kidney disease (ADPKD). ADPKD is primarily characterized by the formation of renal cysts and potential kidney failure. PC1 is an atypical G protein-coupled receptor (GPCR) consisting of 11 transmembrane helices and an autocatalytic GAIN domain that cleaves PC1 into extracellular N-terminal (NTF) and membrane-embedded C-terminal (CTF) fragments. Recently, signaling activation of the PC1 CTF was shown to be regulated by a stalk tethered agonist (TA), a distinct mechanism observed in the adhesion GPCR family. A novel allosteric activation pathway was elucidated for the PC1 CTF through a combination of Gaussian accelerated molecular dynamics (GaMD), mutagenesis and cellular signaling experiments. Here, we show that synthetic, soluble peptides with 7 to 21 residues derived from the stalk TA, in particular, peptides including the first 9 residues (p9), 17 residues (p17) and 21 residues (p21) exhibited the ability to re-activate signaling by a stalkless PC1 CTF mutant in cellular assays. To reveal molecular mechanisms of stalk peptide-mediated signaling activation, we have applied a novel Peptide GaMD (Pep-GaMD) algorithm to elucidate binding conformations of selected stalk peptide agonists p9, p17 and p21 to the stalkless PC1 CTF. The simulations revealed multiple specific binding regions of the stalk peptide agonists to the PC1 protein including an “intermediate” bound yet inactive state. Our Pep-GaMD simulation findings were consistent with the cellular assay experimental data. Binding of peptide agonists to the TOP domain of PC1 induced close TOP-putative pore loop interactions, a characteristic feature of the PC1 CTF signaling activation mechanism. Using sequence covariation analysis of PC1 homologs, we further showed that the peptide binding regions were consistent with covarying residue pairs identified between the TOP domain and the stalk TA. Therefore, structural dynamic insights into the mechanisms of PC1 activation by stalk-derived peptide agonists have enabled an in-depth understanding of PC1 signaling. They will form a foundation for development of PC1 as a therapeutic target for the treatment of ADPKD.

## INTRODUCTION

Polycystin-1 (PC1) is the protein product of the PKD1 gene that is mutated in the majority of cases (∼85%) of autosomal dominant polycystic kidney disease (ADPKD)^1^. ADPKD is a potentially lethal disease, affecting >0.6 million individuals in the US. It causes renal cyst formation that could consequently lead to kidney failure. Currently, the only approved treatment for ADPKD is Jynarque^TM^, a small-molecule antagonist of the arginine vasopressin receptor 2, V2R, whose signaling, and production of cAMP has been shown to be increased in PKD. This drug targets one of the aberrant pathways downstream from the PKD gene mutation but is inadequate due to its limitations in only slowing disease progression and causing adverse side effects^2^. ADPKD severity is dependent on the functional level of PC1, and as such, therapies designed to increase the level of PC1 protein, and its functionality are currently being pursued^3–7^. Approximately one-third of PKD1 mutations are non-truncating and could encode partially functional PC1 protein^3, 8–10^. As such, therapeutic treatments that directly target and activate PC1 may represent a promising approach for the treatment of ADPKD. However, this approach remains difficult due to incomplete knowledge of the proximal-most functions of PC1.

PC1 shares characteristics with the Adhesion class of GPCRs (ADGRs), including a conserved GPCR autoproteolysis inducing (GAIN) domain that directs autocatalytic cleavage at an embedded GPCR proteolysis site (GPS) motif^11^. Intramolecular cleavage at the GPS motif generates two non-covalently attached fragments - the extracellular N-terminal fragment (NTF) and the membrane-embedded C-terminal fragment (CTF)^12, 13^. Similar to ADGRs, the PC1 NTF consists of multiple adhesive domains that promote interactions between cells and with the extracellular matrix^14–18^, while the PC1 CTF is composed of 11 transmembrane (TM) helices and a short C-terminal tail (C-tail)^19^ that has been shown to interact with G proteins for signaling activation or regulation^20, 21^ and has thus led to description of PC1 as an atypical GPCR. Previous studies demonstrated the critical importance of cleavage at the PC1 GPS site to prevent renal cystogenesis in mouse models^4, 22^. For the ADGRs, a tethered agonist (TA) model has been proposed for activation of G protein signaling. After dissociation of the NTF, the N-terminal stalk of the ADGR CTF interacts with its membrane-embedded TM domains to induce conformational rearrangements that mediate activation of G protein signaling^23–26^. Exogenous synthetic peptides consisting of various lengths of the N-terminal sequence of the stalk have been shown to function as soluble agonists in activation of signaling by full-length and CTF mutants for numerous ADGRs^20, 27^.

In previous studies of the PC1 CTF, we revealed a stalk TA-dependent molecular mechanism underlying CTF-mediated activation of an NFAT promoter luciferase reporter through complementary *in vitro* cell signaling experiments and all-atom Gaussian accelerated Molecular Dynamics (GaMD) simulations^28^. GaMD is an unconstrained enhanced sampling method that works by adding a harmonic boost potential to reduce large biomolecular energy barriers^29^ and has been used successfully to capture multiple complex biological processes^30–37,38–44,30,32^ including GPCR activation^31^. Expression constructs encoding a stalkless PC1 CTF (a nonbiological mutant with deletion of the first 21 N-terminal residues of CTF) and three ADPKD-associated missense mutants within the stalk region (G3052R, R3063C, and R3063P) were shown to be defective in reporter activation as compared to wildtype PC1 CTF. GaMD simulations revealed a novel allosteric transduction pathway for activation of PC1 CTF signaling that involves initiation by the Stalk interacting with a large extracellular loop between TM segments S1/TM6 and S2/TM7, called the TOP domain, followed by close interactions between the TOP and a putative pore loop (PL) domain between the final 2 TM domains. GaMD simulations of the wildtype PC1 CTF also identified a “Closed/Active” low-energy state related to the large number of Stalk-TOP contacts and the R3848-E4078 ionic interaction between the TOP and PL domains that was not present in the stalkless CTF^28^.

Here, we have utilized *in vitro* cell signaling assays to identify peptide agonists targeting PC1 in combination with *in silico* studies to investigate their binding mechanisms for activation of PC1 signaling. Synthetic peptides of 7-21 residues in length derived from the N-terminus of the PC1 CTF stalk sequence were tested for their ability to re-activate signaling of the stalkless CTF expression construct. Peptide docking and simulations with the recently developed Peptide GaMD (Pep-GaMD), which is able to characterize peptide-protein binding processes more efficiently^45^, were combined for selected peptide agonists p9, p17 and p21 to gain insight into their binding mechanism to the stalkless PC1 CTF. Pep-GaMD was able to successfully refine the docking conformations of the peptides bound to the extracellular TOP domain of PC1. In further Pep-GaMD simulations, the key salt bridge interaction between R3848 and E4078 from the TOP domain and PL, respectively, was observed upon binding of the peptides to stalkless PC1 CTF. Using Potts covariation analysis, in which a protein fitness model is inferred based on observed mutational covariation patterns in multiple sequence alignments (MSAs) of homologous proteins^46^, we identified residues in the PC1 stalk with direct mutational covariation with residues in the TOP domain, which were strikingly consistent with the binding interfaces identified in docking and simulation studies. Overall, these analyses yielded mechanistic insights underlying the stalk peptide agonist mediated signal re-activation of stalkless PC1 CTF. Such insights provide significant contributions toward the future design and development of peptide modulators targeting PC1 for an effective ADPKD therapeutic treatment.

## RESULTS

### Synthetic, stalk-derived peptides re-activate NFAT reporter by CTF^Δst^ *in trans*

Our previous study utilized expression constructs of human PC1 CTF. However, in order to prepare for eventual *in vivo* experiments in mouse models, we generated expression constructs of mouse (m) PC1 consisting of the signal peptide sequence of the T cell surface glycoprotein CD5 (MPMGSLQPLATLYLLGMLVASVLG) fused in frame with the stalk sequence of wildtype mCTF beginning with residue T3041, or with a ‘stalkless’ CTF lacking the first 21 residues of the stalk (mCTF^Δst^) beginning with residue S3062 (**Fig. 1A**). The CD5 signal peptide coding sequence was added to the wildtype mCTF and stalkless mCTF^Δst^ expression constructs in order to ensure their translation at the endoplasmic reticulum for plasma membrane localization.

**Figure 1.**
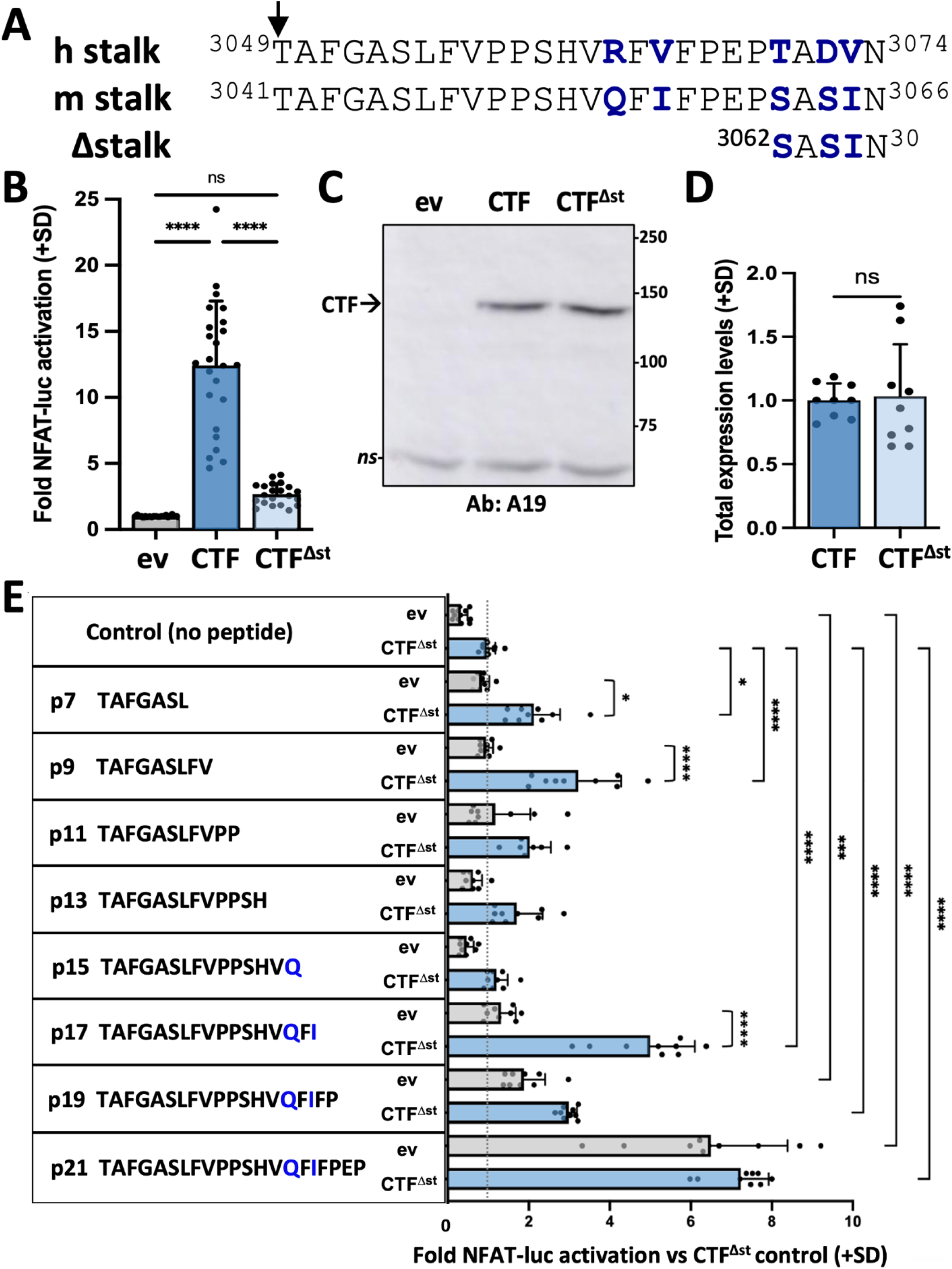
Synthetic peptides derived from the stalk sequence of PC1 can stimulate signaling of stalkless PC1 CTF. (**A**) Alignment of CTF stalk sequences from human (h) and mouse (m) PC1. CTF^Δst^ has a 21-residue deletion from the N-terminal end of the stalk region. Arrow, GPS cleavage site. Non-identical residues shown in bolded blue. **(B)** Activation of the NFAT-luc reporter by transfected mCTF or mCTF^Δst^ expression constructs shown relative to empty expression vector (ev) as means (+ standard deviation, SD) of 3 wells/construct from each of 7 independent experiments. (**C**) Representative Western blot of total cell lysates from one of the experiments in (B), probed with antisera A19 against mouse PC1 C-tail. ns, non-specific. (**D**) Summary of the total expression levels (means +SD) of CTF^Δst^ relative to CTF from the experiments in (B). (**E**) Stalk peptide treatment of ev-or mCTF^Δst^-transfected cells. Sequences of stalk-derived peptides p7-p21 are shown. Graph represents the fold NFAT-luc activation for both eV- (gray bars) and CTF^Δst^- (blue bars) transfected cells relative to the CTF^Δst^ control after 24 hr treatment with or without peptide. Results are the means (+SD) of 3 separate experiments, each with 3 wells/condition. *, p < 0.05; ***, p = 0.0001; ****, p < 0.0001. Analysis by 1-way ANOVA with Tukey-Kramer post-test.

Transient transfection of HEK293T cells with either empty expression vector (ev), CTF or CTF^Δst^ showed that the CTF^Δst^ mutant exhibited a dramatic loss of NFAT reporter activation that was essentially reduced to ev control levels (**Fig. 1B**). Both total (**Fig. 1C-D**) and cell surface (**Fig. S1A-B**) expression levels of CTF^Δst^ were comparable to CTF, which suggests that neither protein stability nor membrane trafficking was responsible for the inability of CTF^Δst^ to activate the NFAT reporter. These results are consistent with those obtained using expression constructs of human PC1 that demonstrated the stalk region of PC1 CTF acts as a tethered peptide agonist^28^.

To further investigate the agonistic property of the CTF stalk, we synthesized peptides (p) consisting of the N-terminal 7, 9, 11, 13, 15, 17, 19 or 21 residues from the stalk sequence of mPC1. All peptides were appended with a C-terminal, 7-residue hydrophilic sequence (GGKKKKK) to increase solubility. HEK293T cells were transiently transfected with empty expression vector or mCTF^Δst^ plasmids along with the NFAT luciferase reporter and then treated with stalk peptides p7 through p21 or with addition of culture medium only (‘no peptide’ control).

The NFAT reporter was significantly activated in CTF^Δst^-transfected cells by treatment with p7, p9 or p17 as compared to their corresponding ev + peptide treatment controls. These stalk peptides also significantly increased reporter activity in comparison to the CTF^Δst^ with no peptide treatment control (**Fig. 1E**). Treatment of CTF^Δst^-transfected cells with p19 or p21 also significantly increased reporter activation in comparison to the CTF^Δst^ + no peptide control; however, reporter activation occurred in both ev- and CTF^Δst^-transfected cells treated with either p19 or p21, suggesting that p19- and p21-mediated activation was not dependent on exogenous expression of mouse CTF^Δst^ and could be activating the endogenous human PC1 protein. Treatment of ev- and CTF^Δst^-transfected cells with peptide consisting of the hydrophilic sequence alone (i.e., solubility peptide) showed that the solubility tag was not responsible for the rescue of CTF^Δst^-mediated reporter activation by stalk peptides such as p17 (**Fig. S2**). Altogether, these results were consistent with soluble stalk-derived peptides acting as PC1 CTF agonists *in trans*, and provided additional support for the PC1 CTF stalk region harboring tethered agonist (TA) activity^28^. We hypothesized that the soluble, activating peptides bind to the TOP domain of PC1 in a manner mimicking the tethered stalk in order to reactivate signaling of the stalkless CTF^Δst^ mutant. From among the active stalk-derived peptides, we selected p9, p17 and p21 that exhibited the highest agonist activity in CTF^Δst^ reporter activation (**Fig. 1E**) for *in silico* simulation studies.

### Docking and Pep-GaMD simulations of peptide agonist binding to stalkless PC1 CTF

We chose to use the stalkless CTF (ΔStalk CTF) as representing the least complex system in which the binding of exogenous peptides could be studied. ΔStalk CTF is not a biological form or a mutant protein of PC1 observed in ADPKD. However, in our previous study, it mimicked the ADPKD-associated stalk mutants by being defective in cell signaling assays and rarely formed the R3848-E4078 salt bridge that was frequently seen in GaMD simulations with wild type CTF^28^. Therefore ΔStalk CTF served as the negative control for the following studies.

The cryo-EM structure of human PC1-PC2 complex (PDB: 6A70)^47^ was used to build the computational model for ΔStalk PC1 CTF after deleting the first 21 residues (3049-3069) from the CTF^28^. We successfully docked the p9, p17 and p21 stalk peptides to the ΔStalk CTF model with HPEPDOCK^48^ (**See SI**). The peptides all bound to the TOP domain and the interface between the TOP domain and extracellular loop 1 (ECL1) of CTF (**Fig. S3A-B**). In particular, peptide p21 occupied a closely similar binding region as the stalk in wildtype CTF as observed in the previous study^28^. We then performed five independent 500 ns Pep-GaMD simulations on each of the three stalk peptide agonists p9, p17 and p21 bound to ΔStalk CTF to refine their HPEPDOCK docking conformations (**See SI**).

With the Pep-GaMD simulation frames, we performed structural clustering of each peptide using the hierarchical agglomerative algorithm in CPPTRAJ^49^. The top-ranked conformations of each peptide bound to ΔStalk CTF were compared to their initial docking conformations. Next, we calculated 2D free energy profiles of the peptides-bound ΔStalk CTF by reweighting the Pep-GaMD simulations. The R3848-E4078 residue distance and the number of contacts between the peptides (p9, p17 and p21) and TOP domains were selected as the reaction coordinates. The number of contacts was calculated between any atom pairs within 4 Å distance of the peptide and extracellular domains of PC1 protein. In the subsequent analyses, stalk and peptide residues are numbered relative to the N terminus of the stalk as starting from 1, while residues of the Δstalk CTF are numbered according to the human PC1 protein sequence.

### Active conformation of peptide p9-bound PC1 CTF

From the free energy profile of the p9-bound ΔStalk CTF, we identified “Unbound” and “Bound” low-energy states (**Fig. 2A**). In the docking conformation, peptide p9 bound to the interface between the TOP and ECL1 of ΔStalk CTF (**Fig. S3A-B**). In Pep-GaMD simulations, the p9 peptide dissociated from the TOP-ECL1 binding pocket and rebound to the TOP domain in a slightly different region (**Fig. 2B-C**). The p9 peptide sequence is mostly composed of hydrophobic residues. Polar interactions between the main chain atoms of peptide-protein residues were observed in the top-ranked representative conformation of the p9-bound ΔStalk CTF. Protein residues R3892 and H3864 formed hydrogen bonds with p9 residues A2 and A5, respectively (**Fig. 2D**). These interactions were also highlighted in the protein contact map between peptide p9 and the extracellular domains of CTF in the representative “Bound” state **(Fig. S4)**. The distance between the TOP domain residue R3848 and PL residue E4078 was 3.9 Å (**Fig. 2E**), suggesting that the top-ranked representative conformation of the p9-bound ΔStalk CTF was in the “Closed/Active” low-energy state. In addition, the 2D free energy profile of each individual simulation was calculated and the “Bound” low-energy state was consistently identified in the 2D free energy profiles of peptide p9 **(Fig. S5)**.

**Figure 2.**
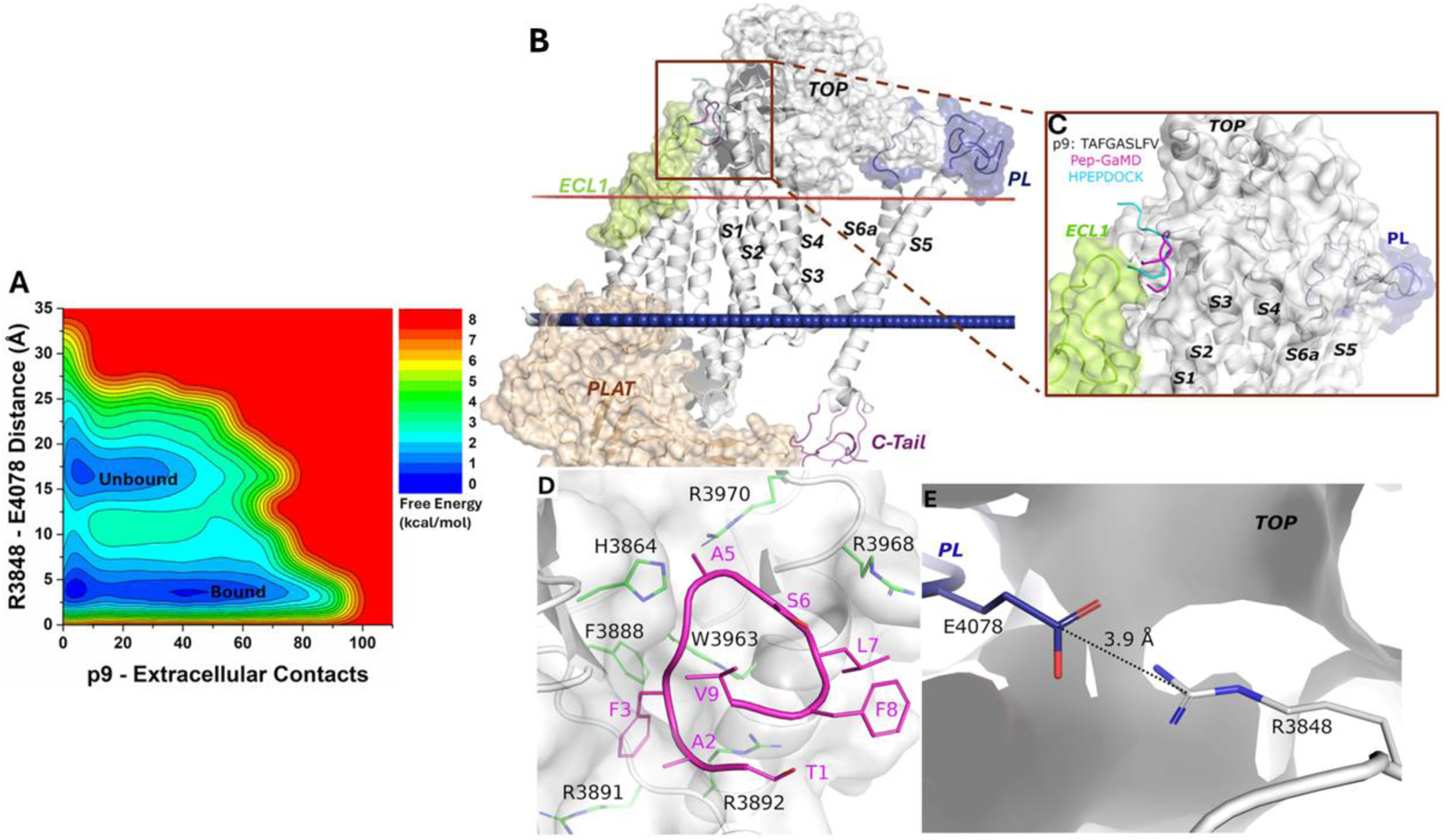
(**A**) Free energy profile of the p9-bound ΔStalk CTF regarding the number of atom contacts between p9 and extracellular domains of CTF and the distance between the CZ atom of R3848 and the CD atom of R4078 in CTF calculated from Pep-GaMD simulations. (**B-C**) Comparison of HPEPDOCK docking (cyan) and Pep-GaMD refined (magenta) conformations of peptide p9 in 𝞓Stalk CTF. (**D**) Polar interactions between peptide-protein residues observed in the top-ranked representative conformations of p9. Peptide residues are numbered relative to the N terminus of the stalk with the peptide starting from 1, while residues within Δstalk CTF are numbered according to the standard PC1 residue number. (**E**) Distance between TOP domain residue R3848 and PL residue E4078 observed in p9-bound ΔStalk CTF.

### Active and Intermediate conformational states of peptide p17-bound PC1 CTF

The free energy profile of the p17-bound ΔStalk system allowed us to identify three low-energy states - “Unbound”, “Intermediate”, and “Bound” (**Fig. 3A**). In the docking conformation, peptide p17 bound to the interface between the TOP and ECL1 of ΔStalk CTF (**Fig. S3A-B**). In the Pep-GaMD refined “Bound” state, a folded antiparallel ß-strand conformation was observed for the peptide p17 at the interface of ECL1 and the TOP domain (**Fig. 3B-C**). Peptide residues T1, F3, A5, F8, F16 and V17 formed hydrophobic interactions with the protein residues H3311, R3314 and Y3307 from ECL1, and E3708, S3711, Q3707, A3704, R3700 and L3701 from the TOP domain (**Fig. 3D**). These interactions were also highlighted in the protein contact map between peptide p17 and the extracellular domains of CTF in the representative “Bound” state **(Fig. S4)**. The distance between the TOP domain residue R3848 and PL residue E4078 was 4.1 Å (**Fig. 3E**), suggesting that the top-ranked representative conformation of the p17 bound ΔStalk CTF was in the “Closed/Active” low-energy state.

**Figure 3.**
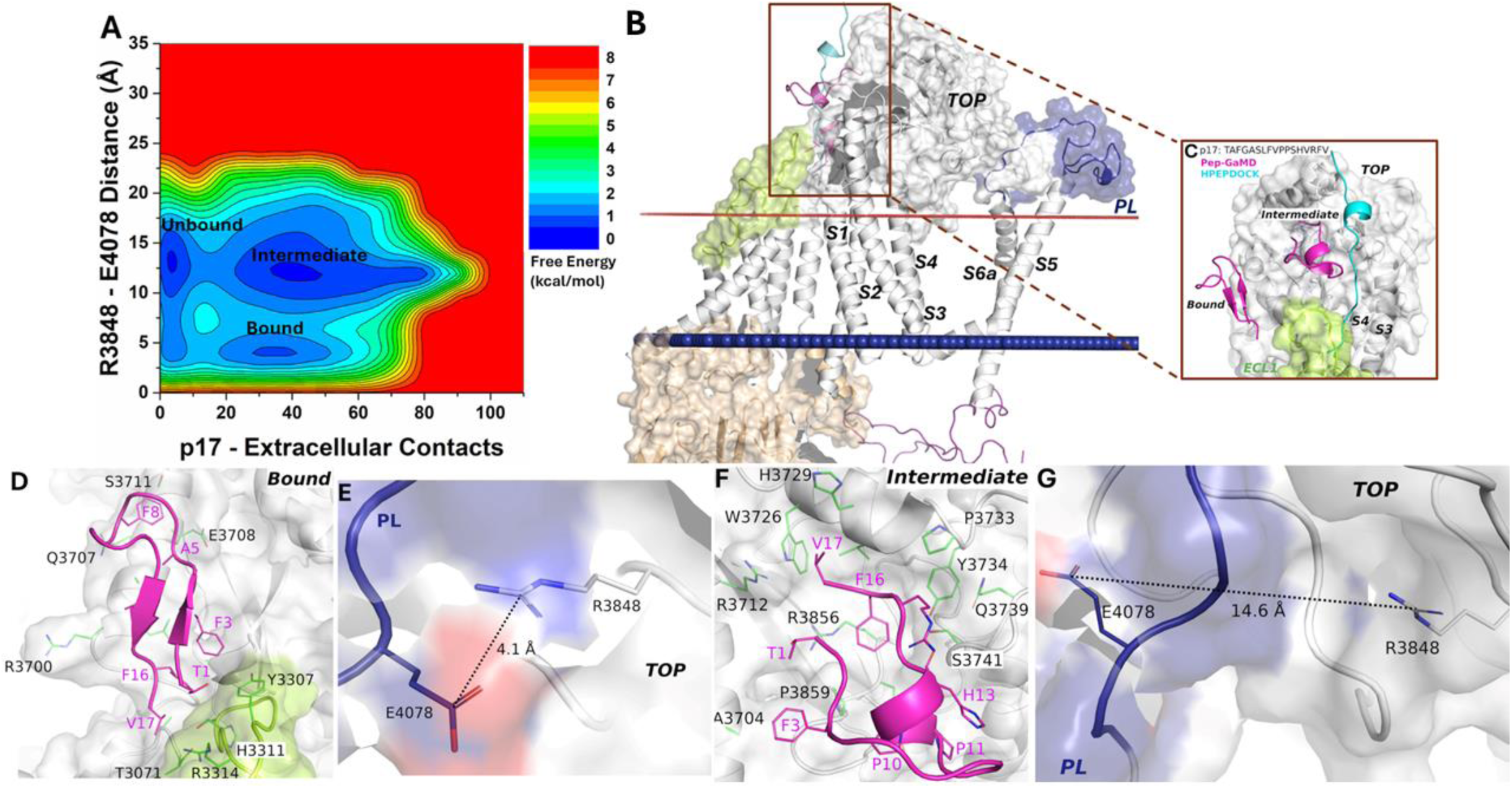
(**A**) Free energy profile of the p17-bound ΔStalk CTF regarding the number of atom contacts between p17 and extracellular domains of CTF and the distance between the CZ atom of R3848 and the CD atom of R4078 in CTF calculated from Pep-GaMD simulations. (**B-C**) Comparison of HPEPDOCK docking (cyan) and Pep-GaMD refined (magenta) conformations of peptide p17 in 𝞓Stalk CTF. Hydrophobic interactions (red dashed lines) between peptide-protein residues observed in the (**D**) “Bound” and (**F**) “Intermediate” low-energy conformations of p17-bound ΔStalk CTF. Distance between TOP domain residue R3848 and PL residue E4078 observed in the **(E)** “Bound” and **(G)** “Intermediate” low-energy conformations of p17-bound ΔStalk CTF.

In the “Intermediate” state, p17 with a short helical turn was also observed to bind the TOP domain of ΔStalk CTF (**Fig. 3B-C**). Hydrophobic residue interactions were also formed between the peptide and protein. In particular, peptide residues T1, F3, P10, P11, H13, R15, F16 and V17 formed hydrophobic interactions with the protein residues P3859, A3704, S3741, Q3739, Y3734, P3733, H3729, W3726, R3712 and R3856 from the TOP domain (**Fig. 3F**). The distance between the TOP domain residue R3848 and PL residue E4078 was 14.6 Å (**Fig. 3G**), suggesting that this representative conformation (ranked the second among the Pep-GaMD structural clusters) of the p17-bound ΔStalk CTF was in the “Intermediate” low-energy state. In addition, the 2D free energy profile of each individual simulation was calculated. Pep-GaMD simulations were able to refine the peptide conformation from the “Unbound” to “Intermediate” and “Bound” states in Sim1 and Sim5, while the peptide reached only the “Intermediate” state in the other three simulations **(Fig. S6)**. The free energy values of 2D PMF minima shown in **Fig. 3A** could differ from those in the 1D PMF minima of peptide structural clusters, especially with the usage of distinct reaction coordinates.

### Active conformational state of peptide p21-bound PC1 CTF

Finally, the free energy profile of the p21-bound ΔStalk CTF allowed us to identify only a broad low-energy well corresponding to the “Bound” state (**Fig. 4A**). The docking conformation of p21-bound ΔStalk CTF was refined through Pep-GaMD simulations, where folding of the peptide was observed on the protein surface of the TOP domain (**Fig. 4B-C**). The p21 peptide occupied a similar binding region as the stalk in wildtype CTF as observed in the previous study^28^. Hydrophobic contacts were observed between peptide residues L7, F8, P10, S12, H13, V14, V17, P19, E20 and P21 and protein residues L3863, L3701, I3705, L3709, E3708, R3712, F3714, H3729, W3726, V3730, L3732, P3733, N3738, R3856 and S3741 (**Fig. 4D**). These interactions were also highlighted in the protein contact map between peptide p21 and the extracellular domains of CTF in the representative “Bound” state **(Fig. S4)**. The distance calculated from the top-ranked structural cluster of the system between the TOP domain residue R3848 and PL residue E4078 was 3.8 Å, corresponding to the “Closed/Active” low-energy state (**Fig. 4E**). Furthermore, time courses of the radius of gyration (*Rg*) and root-mean square deviation (RMSD) of p21 relative to the starting HPEPDOCK conformation showed large conformational changes of the peptide during Pep-GaMD simulations (**Fig. S7)**. In addition, the 2D free energy profile of each individual simulation showed that Pep-GaMD was able to refine the peptide docking conformation to the “Bound” state in all the five simulations **(Fig. S8)**.

**Figure 4.**
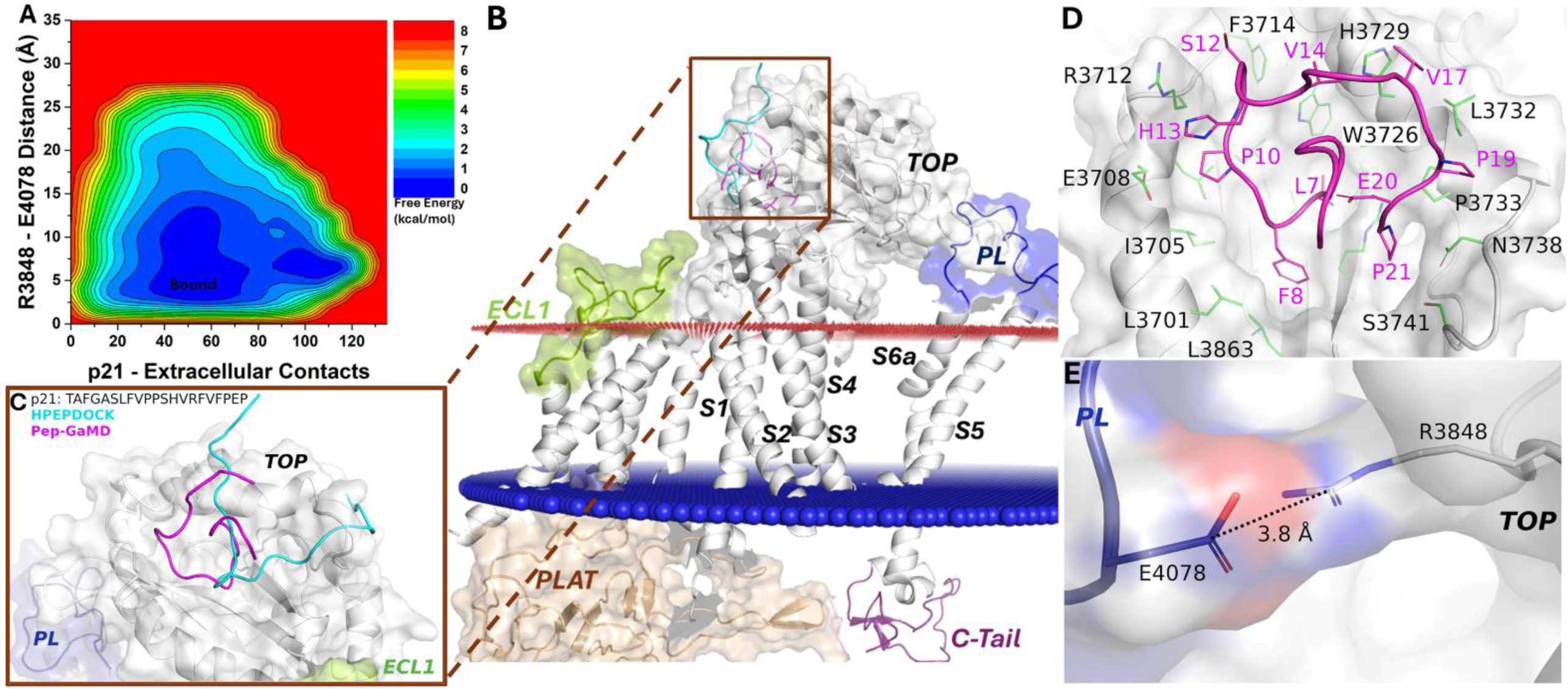
(**A**) Free energy profile of the p21-bound ΔStalk CTF regarding the number of atom contacts between p21 and extracellular domains of CTF and the distance between the CZ atom of R3848 and the CD atom of R4078 in CTF calculated from Pep-GaMD simulations. (**B-C**) Comparison of HPEPDOCK docking (cyan) and Pep-GaMD refined (magenta) conformations of peptide p21 in 𝞓Stalk CTF. (**D**) Polar interactions between peptide-protein residues observed in the top-ranked representative conformations of p21. (**E**) Distance between TOP domain residue R3848 and PL residue E4078 observed in p21-bound ΔStalk CTF.

### Peptide binding regions correlated with covarying residue pairs identified between the TOP domain and stalk TA

To provide an independent basis of evidence supporting the observation of “Bound” and “Intermediate” states of agonist peptide binding, we constructed a multiple sequence alignment (MSA) with an effective count of 1022 evolutionarily diverged PC1 homologs (illustrated in **Fig. S9)** from which we inferred a Potts statistical model (**Fig. 5**). Columns of the MSA with “direct” statistical interactions, as detected using the Potts inference method, reflect compensatory mutation pairs maintained through evolution supporting a conserved function. We limited our MSA to 394 residues on the extracellular side of PC1 because of the computational challenge of fitting the entire PC1 sequence (**Fig. 5B** and **Fig. S9**).

**Figure 5.**
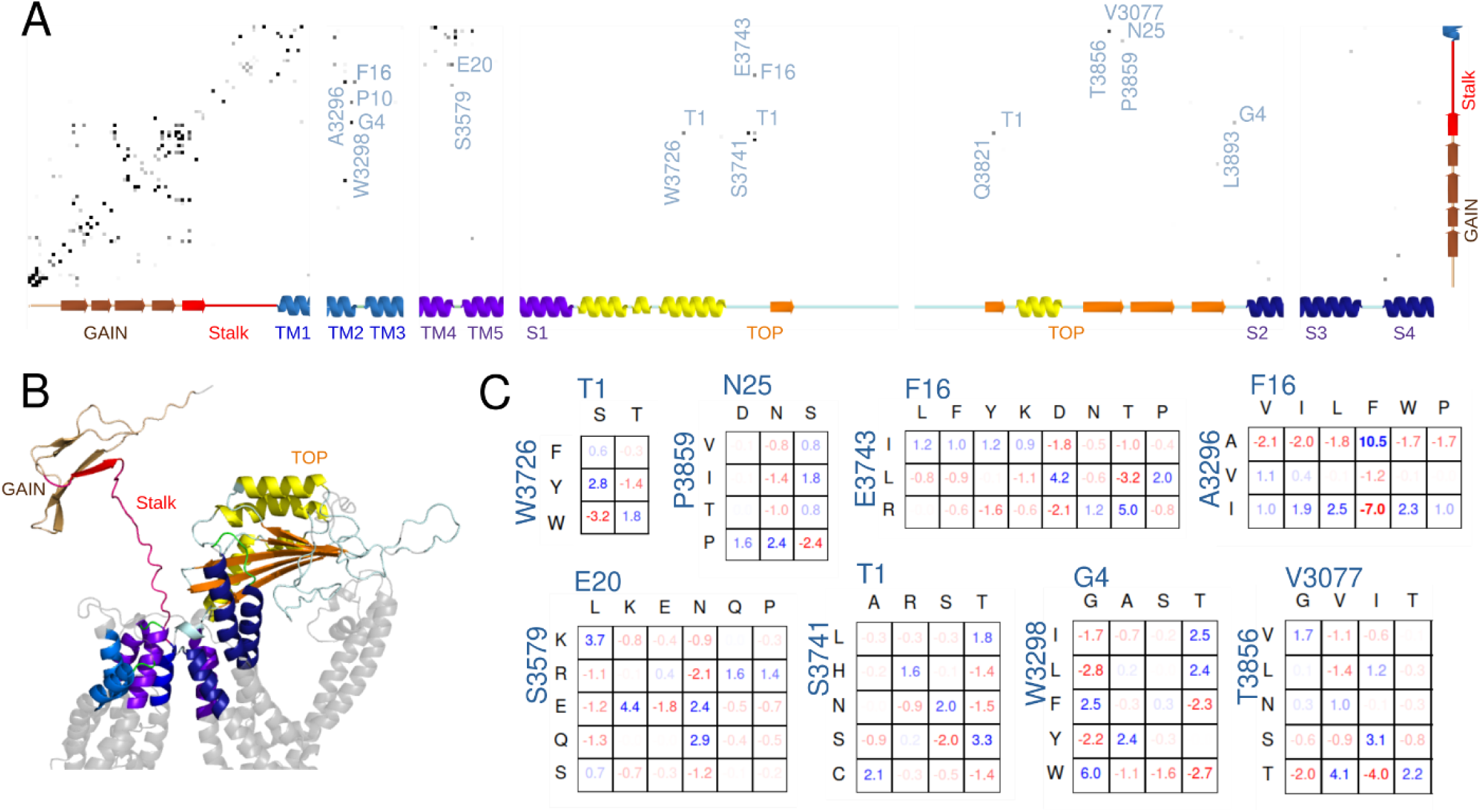
(**A**) Potts interaction map based on the PKD1 multiple-sequence-alignment illustrated in **Figure S4**, showing interactions with the stalk. Gray dots are shown for residue position-pairs with Potts covariation scores above a threshold, colored darker for higher scores, and selected interacting pairs are annotated with the stalk residue (horizontal, numbered from the stalk N-terminus) and other residue (vertical, standard PKD1 numbering) with the PKD1 residue at each position. The secondary structure as a function of position is annotated along the axes. (**B**) Cartoon showing the subset of PC1 included in the Potts covariation analysis colored as in the secondary structure in panel A, using a structure predicted by AlphaFold. Gray regions were excluded from the Potts model. (**C**) Residue Covariation scores for selected position-pairs. The scores reflect the percentage excess frequency of the residue-pair relative to the null expected frequency if the MSA columns were uncorrelated, with blue values reflecting excess and red dearth. Only the most common residue types are shown.

**Fig. 5A** shows the pairs of positions with strong Potts interaction scores where one position is either in the stalk, the nearby GAIN domain, or the TM1 helix. Some predicted interaction pairs recapitulated beta-sheet contacts within the GAIN domain observed in the homolog rat latrophilin-1^11^ as well as predicted by Alphafold^50^ (**Fig. S10**) or between the extracellular ends of the TM2/4 and TM5/6 alpha helices known from cryo-EM structures^47^ or predicted by Alphafold, validating that our model detected biologically functional interactions.

We identified strong interactions between the stalk and other residues from the Potts model. They were not observed in the cryo-EM structure, in which the flexible stalk is missing. For interactions with the TOP domain, out of the 4875 possible pairs (25 stalk residues by 195 TOP domain residues in our Potts model), this analysis detected a stringent set of 6 strongly interacting pairs. Remarkably, multiple positions in this small set were among those relevant to the “Intermediate” binding conformation of p17 and “Bound” conformation of p21 as identified from the Pep-GaMD simulations. These were W3726 and S3741 in the TOP domain, both interacting with T1 of the stalk, and P3859 interacting with N25 at the end of the stalk **(Fig. 5A)**. Additionally, we identified E3743 to be strongly interacting with F16 in the stalk, and it was also near the observed binding region in the TOP domain for the peptide p17 in the “Intermediate” state near S3741 (**Fig. 3F**). The remaining two strong interactions between the stalk and TOP domain involved Q3821 with T1 and L3893 with G4. L3893 is adjacent to R3892 that was identified to interact with the peptide p9 in the “Bound” state (**Fig. 2D**) and mutating it may affect its neighbor’s positioning. Besides the interactions between the TOP domain and the stalk TA, we also found a set of interactions between the stalk and the extracellular ends of TM2-TM3 helices and TM4-

TM5 helices, in which stalk residues G4, P10, F16 and E20 interact with W3298, A3296, and S3579, respectively, as well as a strong interaction between V3077, three residues past the end of the stalk, and T3856 in the TOP domain. TOP domain residue T3856 was also identified as relevant to the binding region of peptide p17 in the “Intermediate” state (**Fig. 3F)** and peptide p21 in the “Bound” state (**Fig. 4D**). These interactions could additionally play a role in stalk-TA activation or could be related to other functionality such as cleavage in the GPS motif.

To gain further insight and to validate that these detected “direct” interactions reflect biologically meaningful functional interactions and are not artifacts of the data, we examined the residue-specific covariation observed in the MSA (**Fig. 5C**), which measures the difference between the observed pairwise residue frequency and its null expectation under assumption of independent variation. Values greater than ∼1% are commonly found to be indications of a statistically reliable mutational covariation (see **SI Computational Methods**), and many of the covarying pairs discovered between the stalk and TOP domain were significantly above this value. We validated that the covarying residue pairings were consistent with biophysical interaction. For example, for the position-pair 20-3579, here annotated such that the first index is the stalk residue numbered relative to the N terminus of the stalk and the second index is the TOP domain residue numbered according to the human PC1 protein sequence, there were excess residue-pair counts in the MSA consistent with opposite-charge or polar pairing such as K20-E3579, N20-Q3579, and others, and a dearth of repulsive like-charge pairs such as E20-E3579. Similarly, position-pair 1-3741 favored certain combinations of polar residues such as T1-S3741. Other position-pairs appeared consistent with hydrophobic packing interactions, such as F16-A3296, G4-W3298, and T1-W3726. A large residue F or W at position 3298 in the TOP domain was commonly paired with a G at stalk position 4, while a smaller I or L residue at position 3298 was more commonly paired with T at stalk position 4.

Molecular Mechanics/Poisson-Boltzmann Surface Area (MM/PBSA)^51^ analysis was further performed to calculate the binding free energies of peptides p9, p17 and p21 to PC1 CTF and decompose the residue-wise energy contributions using the gmx_MMPBSA software^52^. The relative rank of the overall peptide binding free energies **(Table S1)** was consistent with the experimental signaling data, i.e., p21>p9>p17, for which p21 showed the largest binding free energy value of binding (−40.29±6.94 kcal/mol). Furthermore, we performed residue-wise energy decomposition analysis with MM/PBSA using gmx_MMPBSA software^52^, which allowed us to identify key residues that contributed the most to the peptide binding energies. These included residues T1 and V9 in p9 **(Table S2)**, residues T1, R15 and V17 in p17 **(Table S3)**, and residues P10, P11, P19 and P21 in p21 and residue W3726 in the PC1 CTF **(Table S4)**. The energetic contributions of these residues apparently correlated to the sequence coevolution predicted from the Potts model.

## DISCUSSION

In *in vitro*, cell-based signaling assays, PC1 CTF-mediated activation of the NFAT reporter is dependent on its N-terminal, extracellular stalk, as shown by the loss of reporter activity with the CTF stalk-deletion expression construct, CTF^Δst^ ^28^ (**Fig. 1B**) and by the ability of synthetic, stalk sequence-derived peptides to reactivate signaling by CTF^Δst^ *in trans* (**Fig. 1E**). A series of synthetic peptides derived from the N-terminal sequence of the mouse PC1 CTF stalk were used to determine their agonistic activity in PC1 CTF^Δst^-transfected cells. Notably, treatment with stalk peptides p7, p9, p17, p19 and p21 resulted in significant NFAT reporter activity over CTF^Δst^ control (no peptide treatment), wherein the effects of p7, p9 and p17 were specific to mouse CTF^Δst^-expressing cells. These data are consistent with the stalk peptides acting as soluble TA peptide agonists for PC1 and provide further evidence for the activation of PC1 signaling via an ADGR-like TA mechanism.

In ADPKD, numerous missense mutations reported within the GPCR autoproteolysis-inducing (GAIN) domain that have been shown to prevent or inhibit cleavage at the GPCR-coupled proteolysis site (GPS)^11^. Loss of PC1 GPS cleavage, which is known to cause ADPKD, would likely sequester the stalk tethered agonist within the interior of the GAIN domain, which would presumably interfere with interactions between stalk tethered agonist residues and the remainder of the CTF and thus loss of the stalk-mediated signaling mechanism. Furthermore, there are 10 single nucleotide polymorphisms reported within the stalk sequence (ADPKD Variant Database; https://pkdb.mayo.edu/welcome), most of which were found to significantly reduce CTF-mediated activation of the NFAT reporter^53^. In particular, the ADPKD-associated G3052R stalk mutation that was previously analyzed along with the stalkless CTF by GaMD simulations^28^ has the same reduction in activity as the stalkless CTF in the cellular signaling reporter assays and the same loss of active/closed conformation interactions in GaMD analyses. Therefore, the stalkless CTF was used in our docking and simulation studies as representative of a biologically relevant mutant form of PC1.

To reveal the molecular mechanisms of the soluble stalk-derived peptides, we chose to perform HPEPDOCK docking and novel Pep-GaMD simulations to sample the peptide interactions with the ΔStalk PC1 CTF. Pep-GaMD simulations were able to refine the docking conformations of peptide agonists bound to the ΔStalk PC1 CTF. It is important to note that the free energy profiles calculated from GaMD simulations of PC1 CTF were not fully converged since certain variations were observed among the individual simulations. Nevertheless, these calculations allowed us to identify representative low-energy binding conformations of the peptides. Pep-GaMD simulations sampled an antiparallel ß-strand and a short helical secondary structure of peptide p17 bound to the ΔStalk CTF. Furthermore, peptides p9 and p21 adopted a more folded structure as compared to their disordered loop conformations in the docking poses. We also observed TOP-PL interactions, particularly the salt bridge between residues R3848-E4078 that is a key feature of the stalk TA-mediated activation of signaling for PC1 CTF^28^. Signal transduction was initiated upon binding of the stalk (TA) to the TOP domain, which was transmitted to the PL via a salt bridge formation between residue R3848 in the TOP domain and residue E4078 in the PL. The bound peptide agonists p9, p17 and p21 maintained the ΔStalk CTF in its “Closed/Active” conformation as observed in the wildtype PC1 CTF simulations^28^.

The interacting pairs identified using sequence-based covariation analysis matched the pairs identified by Pep-GaMD simulations, providing complementary evidence of the importance of these interactions and of the existence of the “Bound” and “Intermediate” binding states of the stalk TA and stalk-derived peptide agonist. This suggests that such stalk TA binding states are evolutionarily conserved across PC1 orthologs. Covariation analysis identifies interactions important during any part of the protein lifecycle, and alone cannot be used to distinguish which conformational state an interaction arises in. By comparison to the conformations found in the Pep-GaMD simulations, we found that most of the identified interactions between the stalk TA and TOP domain were consistent with either the “Intermediate” or “Bound” binding states of the stalk-derived peptides, which are related to CTF inactive and active signaling states, however, it remained possible that other interactions, such as between the start of the stalk-TA and TM2/TM3, may be related to conformational states necessary for cleavage of the GAIN/GPS domain. Additionally, structural contacts may be incompletely detected at some positions when the statistical signal of covariation is masked by high conservation, subfamily specialization, or misalignment. This can explain why some interactions identified in the binding interface through docking are not detected using covariation analysis. Despite this, the specific subset of interactions detected using covariation analysis suggest broader peptide binding interfaces, and we found these to be consistent with the peptide binding interactions observed in the Pep-GaMD simulations and the MM/PBSA binding free energy analysis, and the covarying residue pairings were consistent with functional biophysical interactions.

The proposed binding interactions of the PC1 stalk peptides shares some similarity with those observed for the ADGRs. Specifically, Xiao et al. resolved cryo-EM structures of active ADGRG2 and ADGRG4 in complex with tethered Stachel sequences^27^. The structures showed that the 15 residue Stachel sequence inserts into the TM bundle to form intense hydrophobic interactions. A hydrophobic F/Y/LXφφφXφ motif identified in the ADGR tethered sequences formed five finger-like projections in the hydrophobic pits of the TM bundle^27^. In our study, we observed a similar pattern of intense hydrophobic interactions between the peptide agonists p9, p17 and p21 and the hydrophobic pockets in the TOP domain of PC1 CTF. Notably, a closely similar TOP binding pocket was identified for interaction of the tethered agonist (Stalk) in our previous study^28^ and for binding of peptide agonist p21 in this study. The TOP domain hydrophobic pocket may serve as a significant candidate binding site for designing new synthetic peptides or small molecules to aid in rescue of PC1 function levels. Moreover, the shorter peptide agonists’ (p9 and p17) binding sites also serve as novel pockets for design and development of therapeutic approaches for treating ADPKD. While the present study is focused on identification of initial peptides that are able to activate the PC1 CTF, we shall include further mutation experiments and simulations, peptide SAR and optimization of the lead peptides in future studies. It is also important to note that we have not tested the selectivity of the peptides for PC1 versus PC2 in the present study primarily because transfection of PC2 does not activate the NFAT reporter. However, it is possible that co-transfection of PC2 with the PC1 CTF could alter the stalk peptide binding. This will be important to consider in future studies.

## Acknowledgments

We thank Dr. Yan Zhang for her valuable discussions. This work used supercomputing resources with allocation award TG-MCB180049 through the Advanced Cyberinfrastructure Coordination Ecosystem: Services & Support (ACCESS) program, which is supported by National Science Foundation grants #2138259, #2138286, #2138307, #2137603, and #2138296, and project M2874 through the National Energy Research Scientific Computing Center (NERSC), which is a U.S. Department of Energy Office of Science User Facility operated under Contract No. DE-AC02-05CH11231, and the Research Computing Cluster at the University of Kansas. This work was supported in part by National Institutes of Health (R01DK123590 and R56DK135824), Department of Defense CDMRP PRMRP Discovery Award (PR160710/W81XWH-17-1-0301), and Pilot Grant funding from the School of Health Professions at KU Medical Center (to R.L.M.), Pilot award 1004015 from Jared Grantham Kidney Institute at KU Medical Center (to Y.M. and R.L.M.) and startup project 27110 at University of North Carolina - Chapel Hill (to Y.M.), and A.H acknowledges support from National Institutes of Health (R35-GM132090 and OD020095). This research includes calculations carried out on HPC resources supported in part by the National Science Foundation through major research instrumentation grant number 1625061 and by the US Army Research Laboratory under contract number W911NF-16-2-0189.

## Author Contributions

R.M. and Y.M. designed research; S.P., B. S. M., E. N. M. and A.H. performed research; S.P., B. S. M., K.J; E. N. M., A.H., R.M. and Y.M. analyzed data; and S.P., K.J; A.H., R.M. and Y.M. wrote the paper.

## Competing Interest Statement

No competing interests.

## TOC Graphic

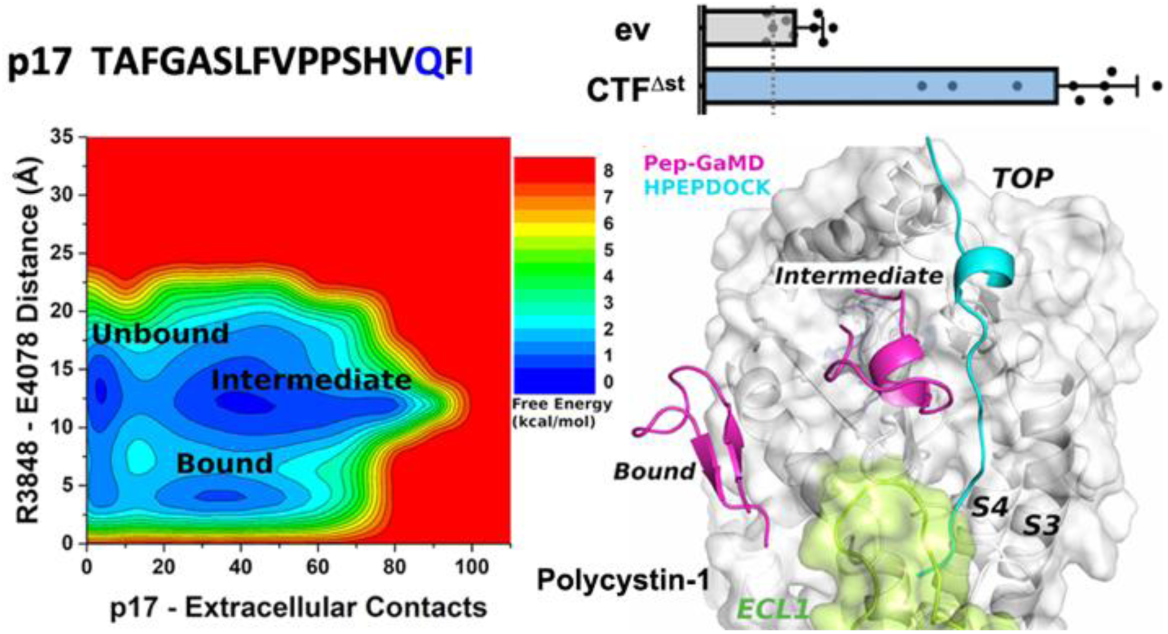
Structural dynamic models are presented for binding of novel synthetic, soluble peptide agonists and associated activation of Polycystin-1 through a combination of complementary cellular signaling assays, accelerated molecular simulations and sequence coevolutionary analysis. The p17 peptide derived from the first 17 residues of the protein stalk tethered agonist is shown here.

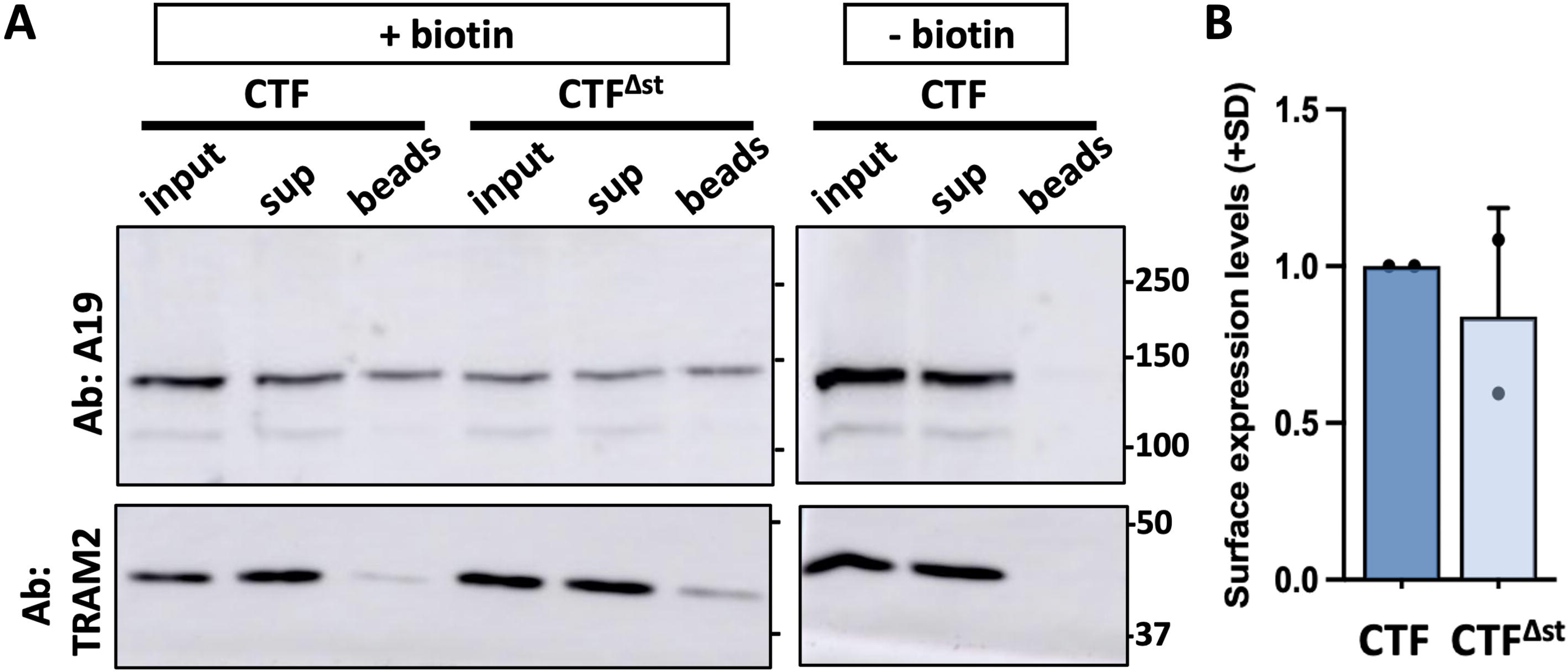

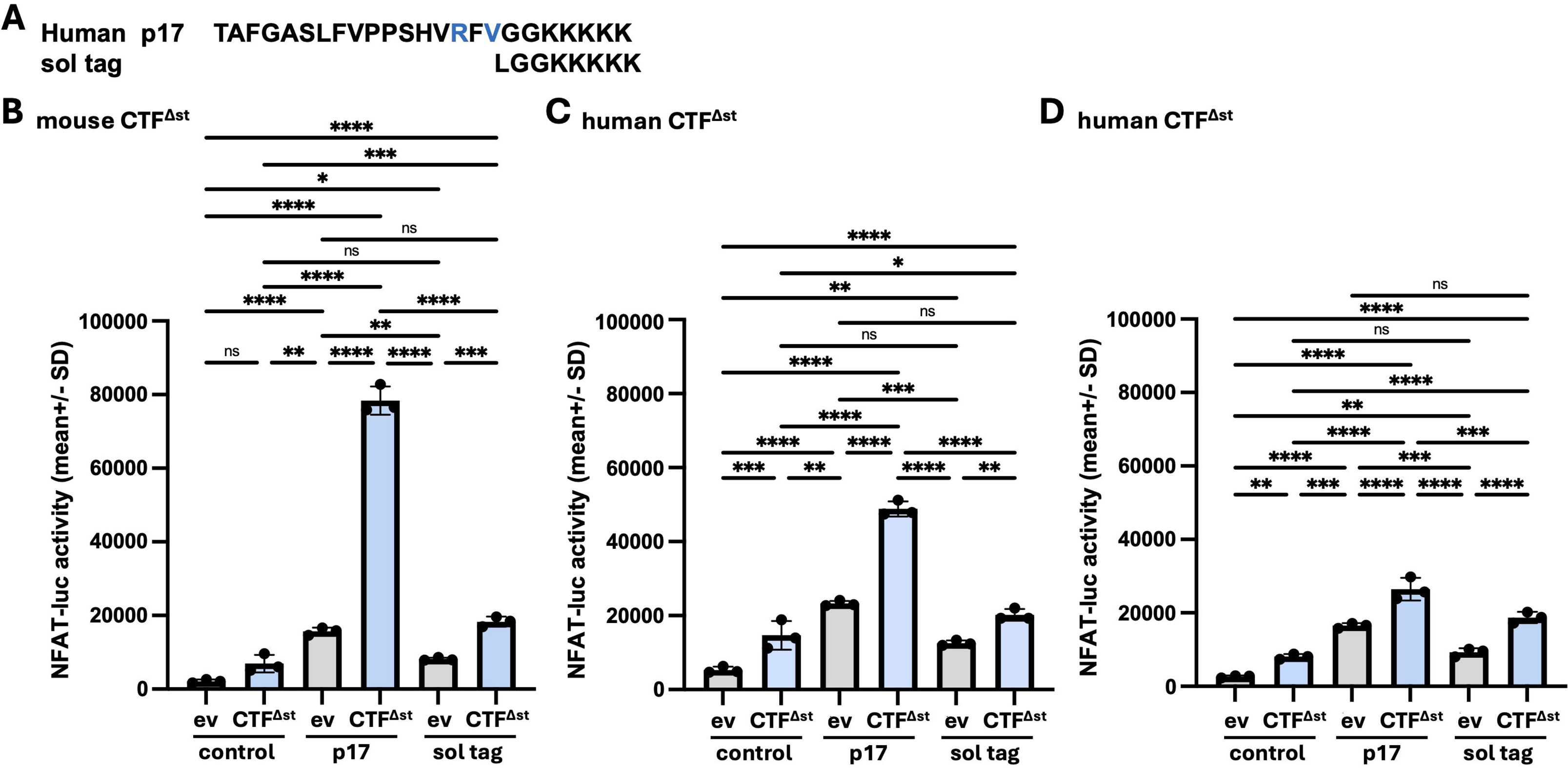

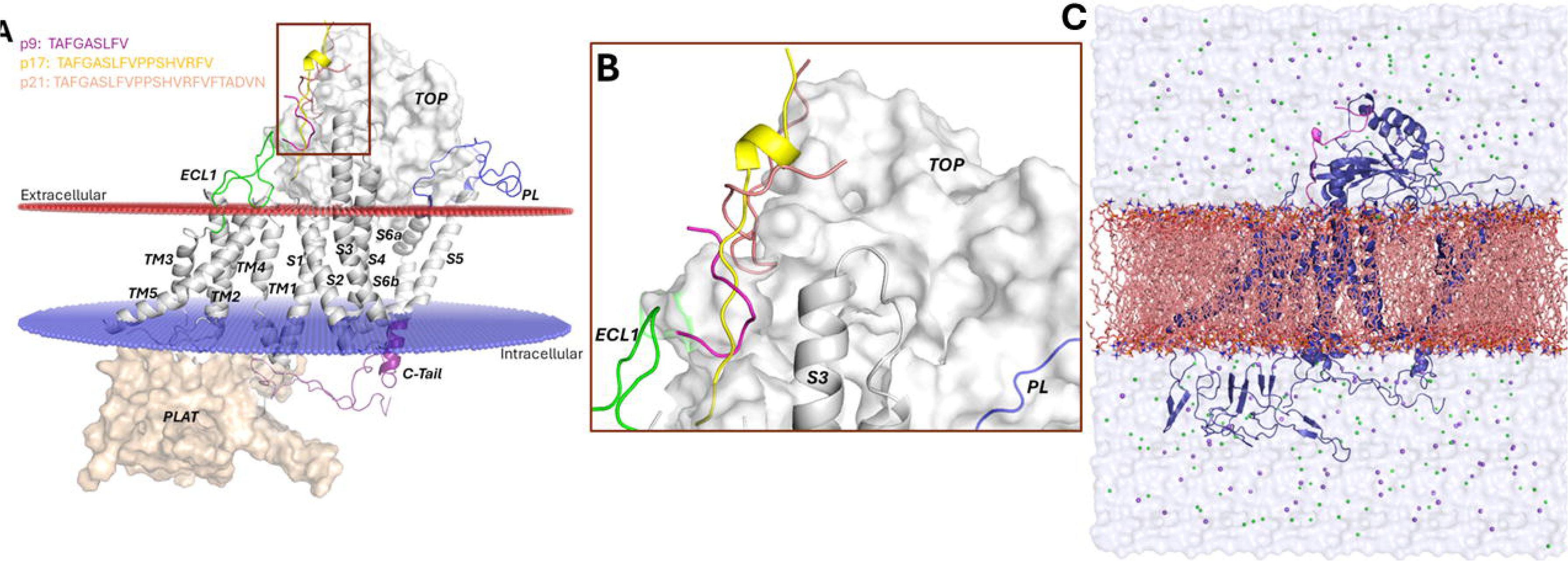

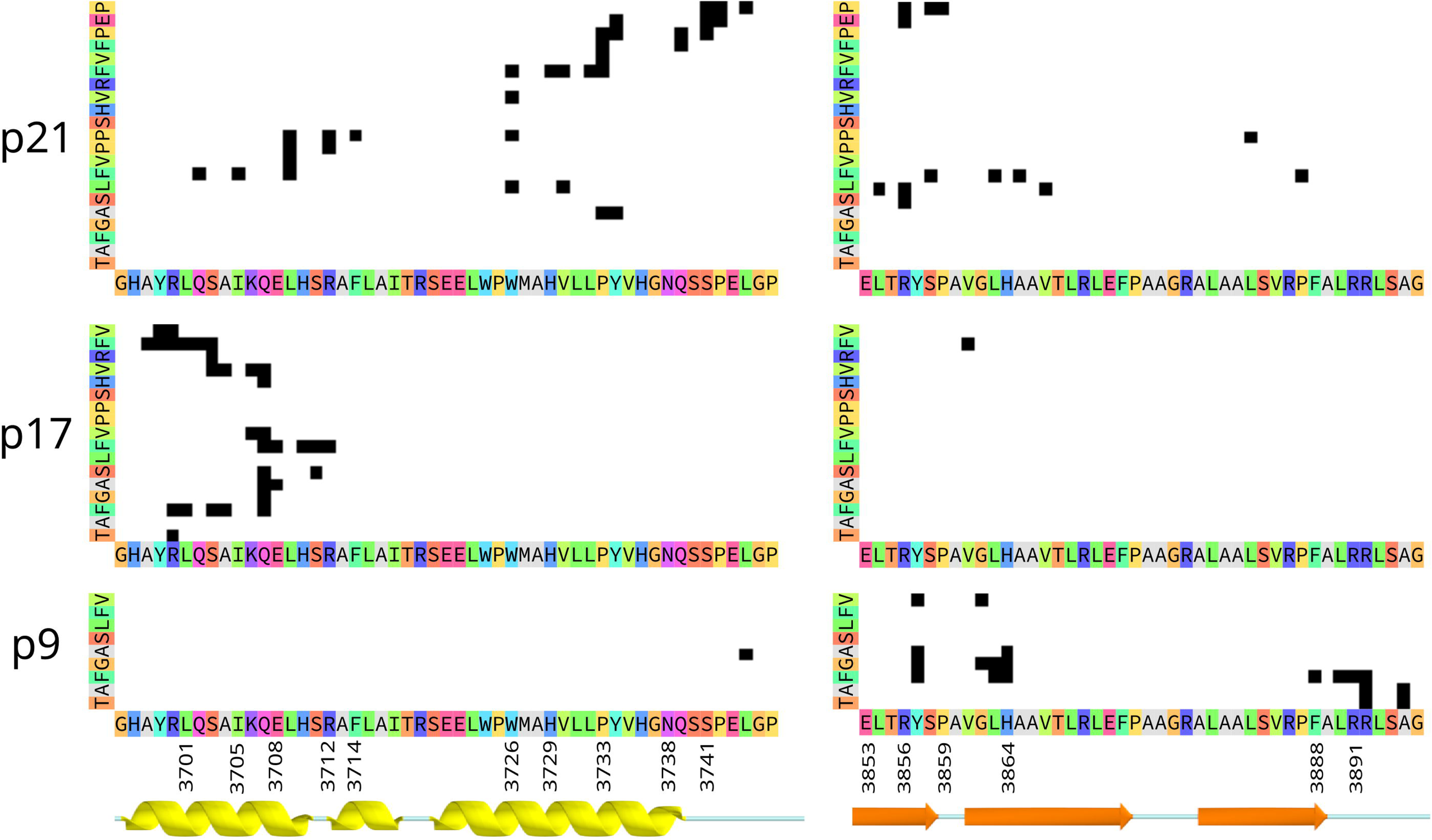

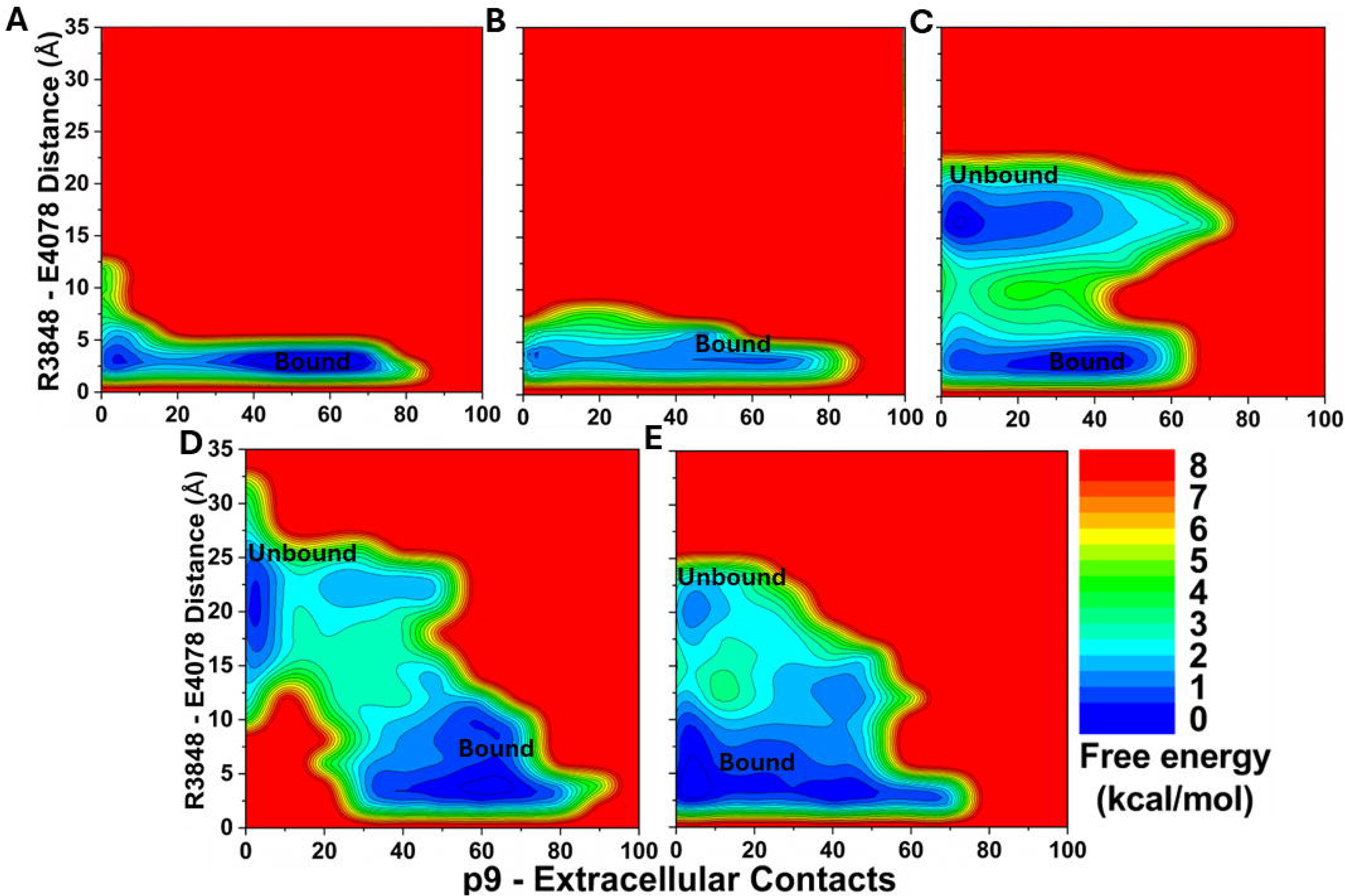

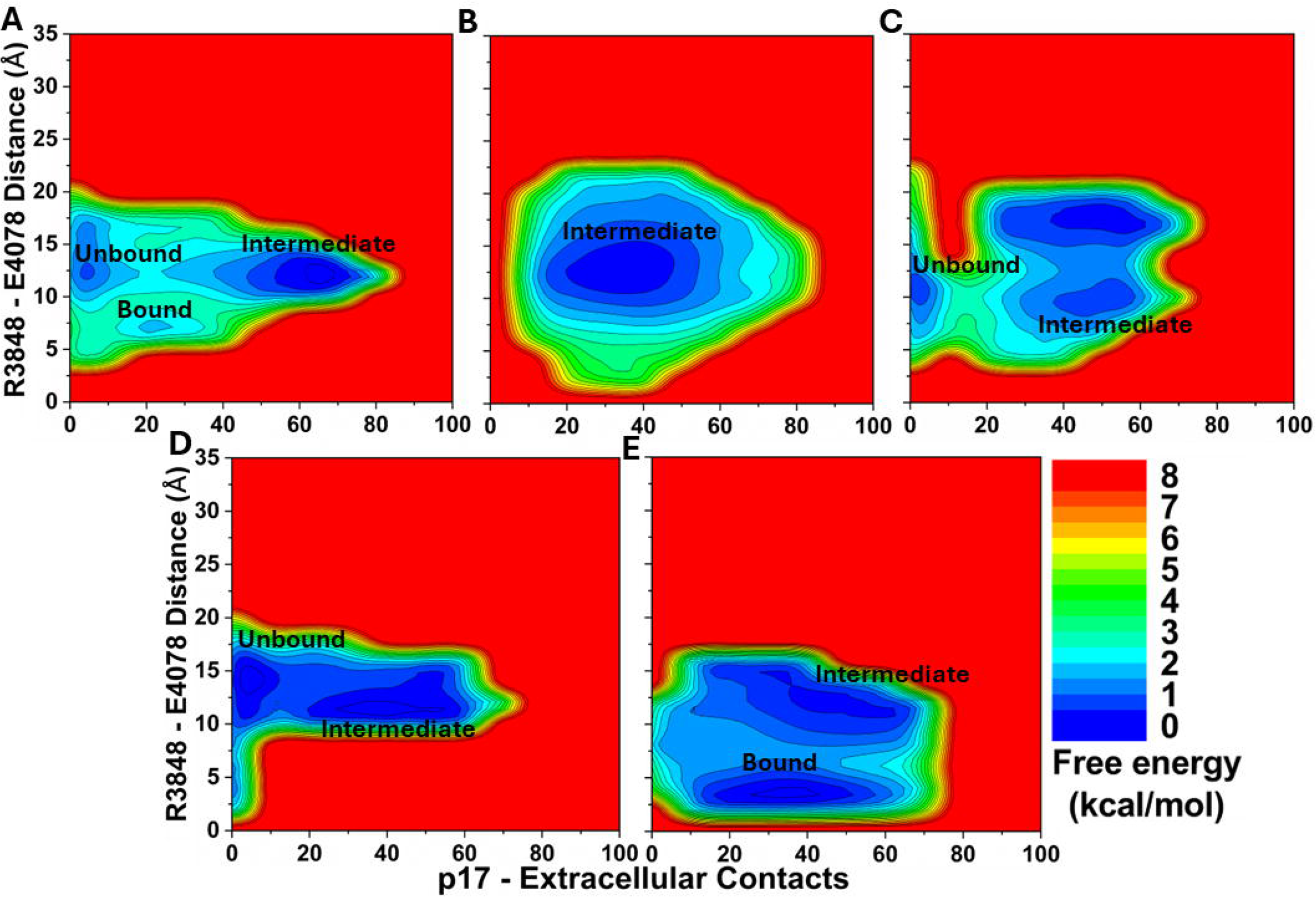

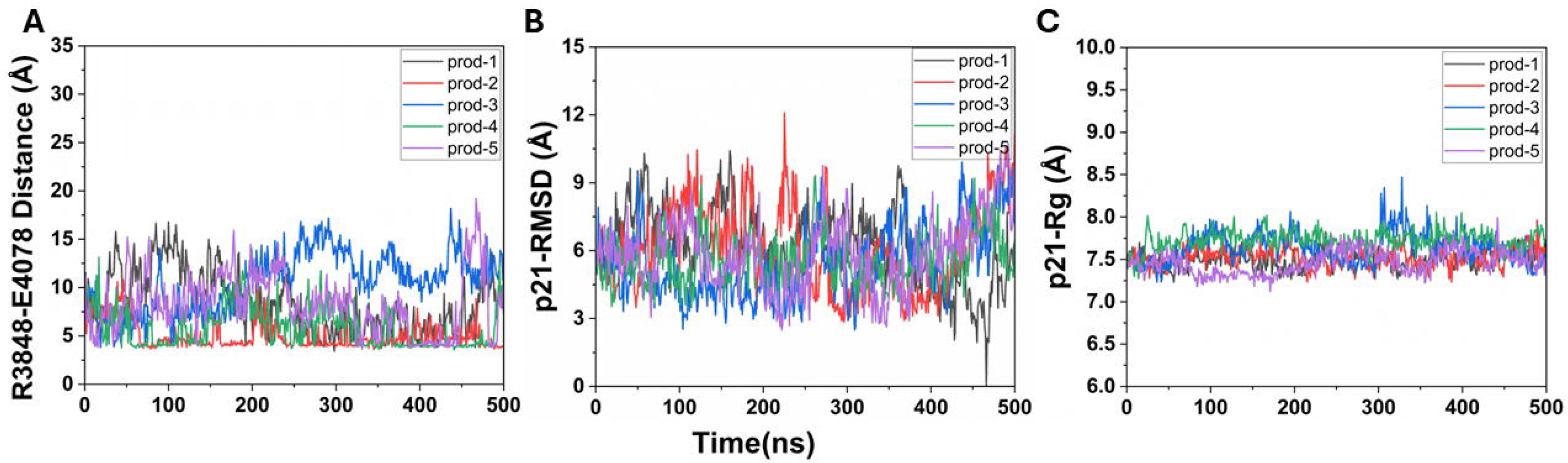

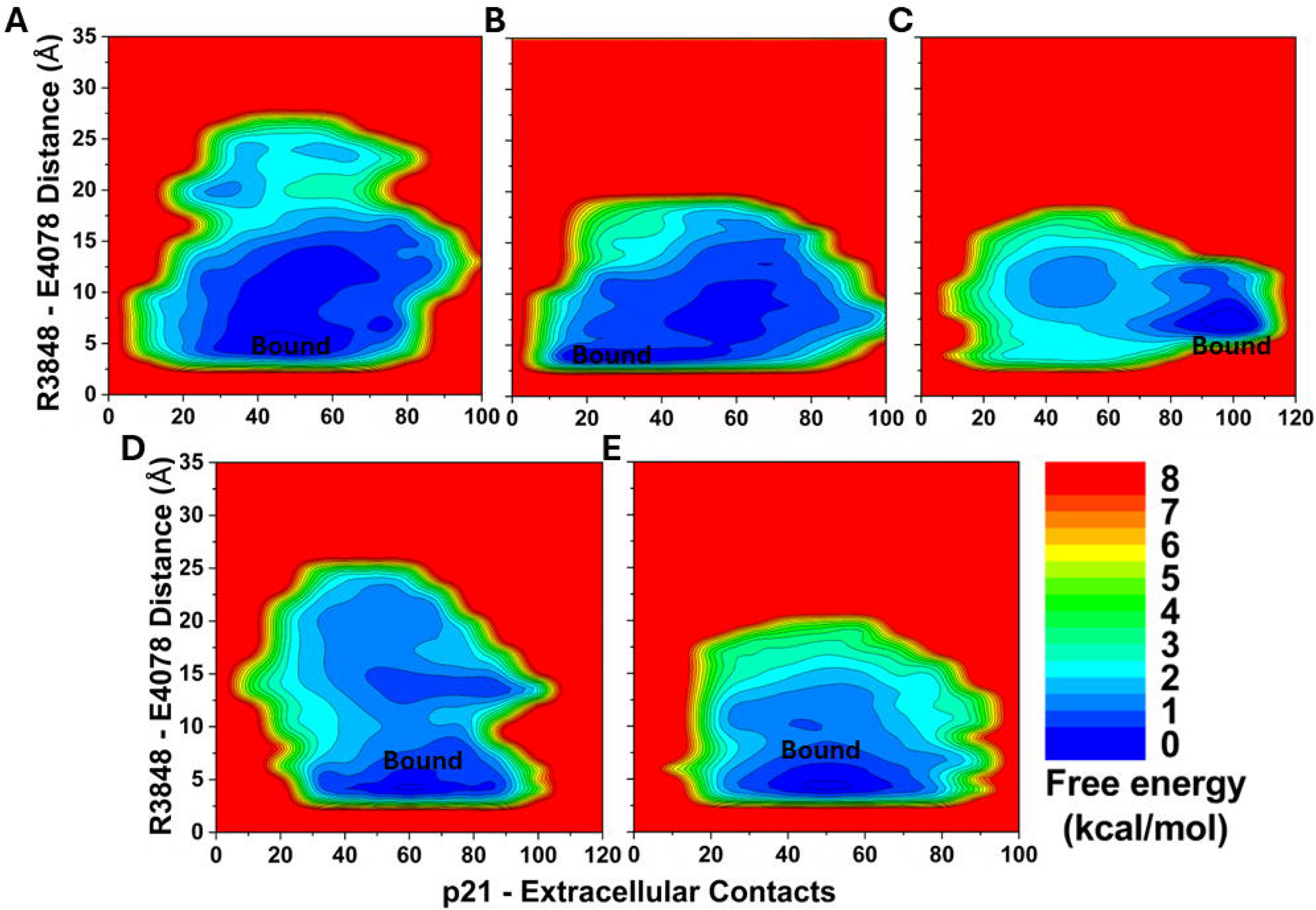

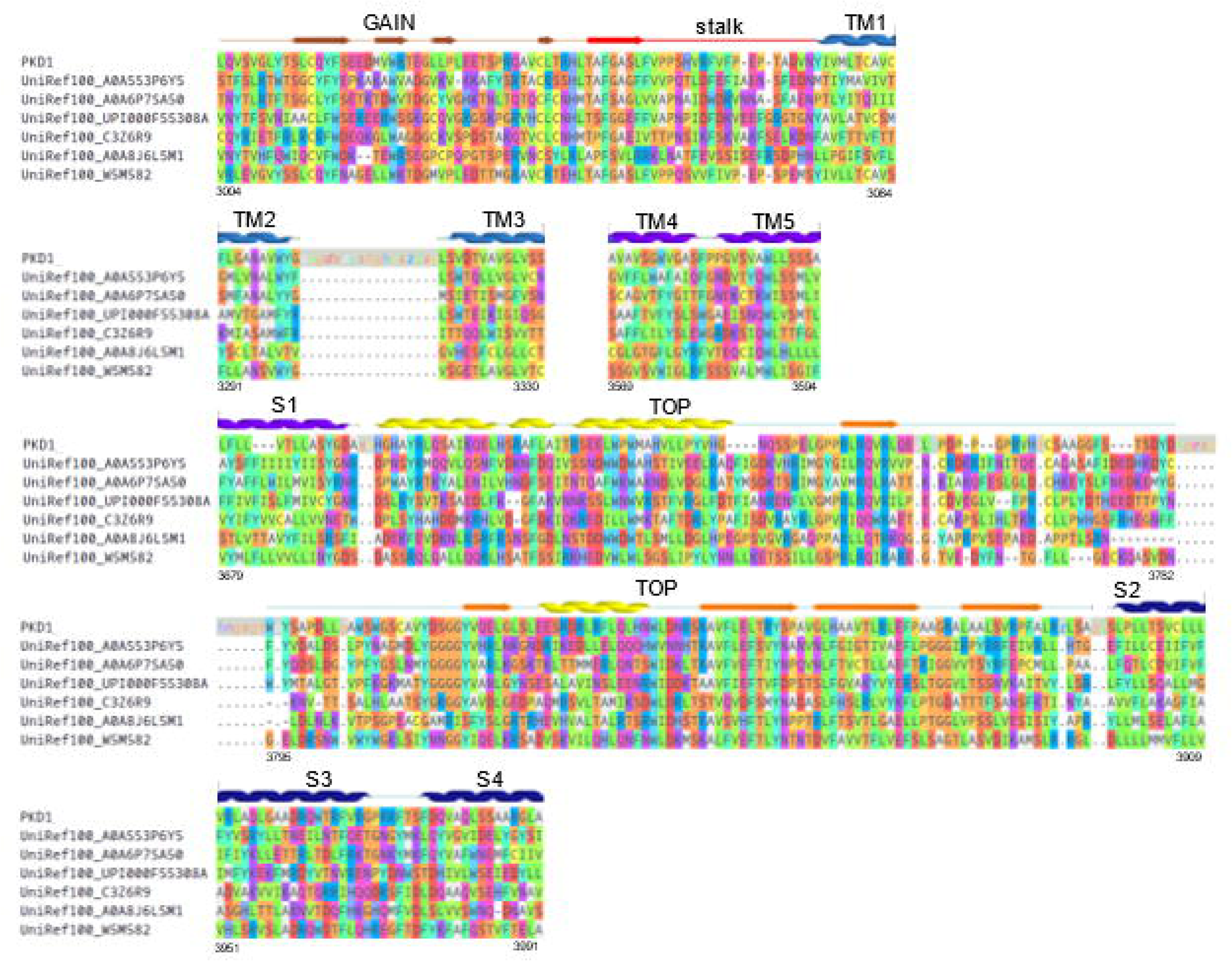

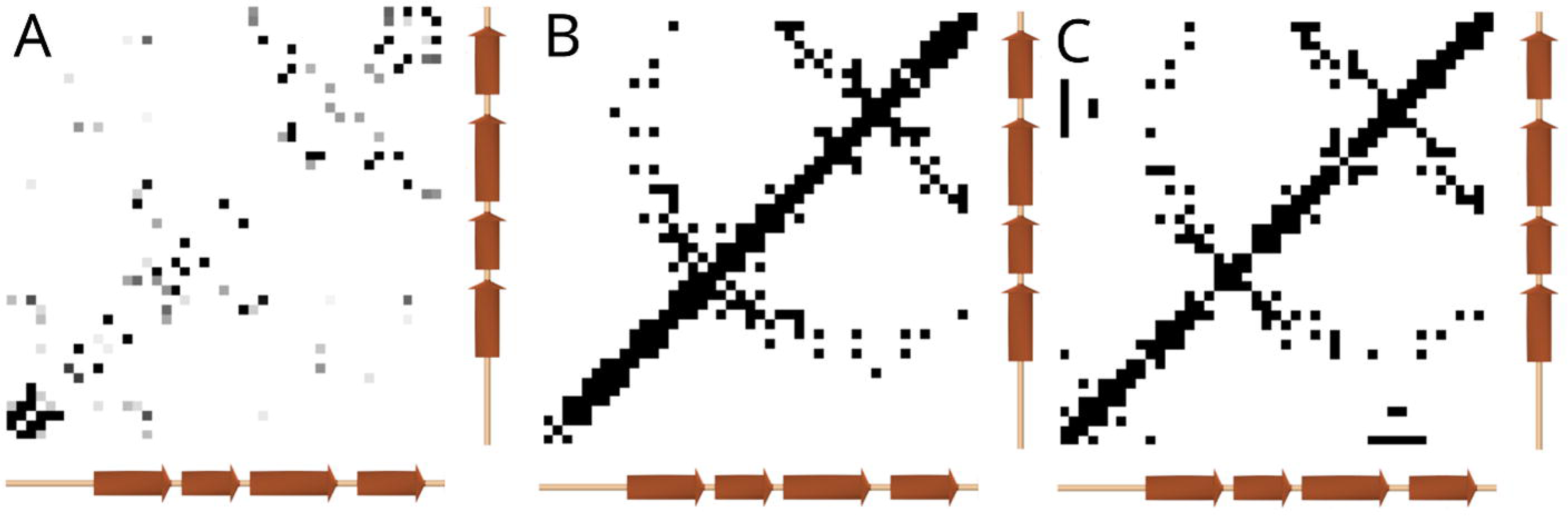

## Notes

### Competing Interest Statement

The authors have declared no competing interest.

### Summary of Updates

1. Study on Stalkless CTF as a control has been added to the revised manuscript. 2. Individual free energy profiles, contact maps and MM/PBSA and energy decomposition analysis has been added for p9 p17 and p21 peptides. 3. The effect of the hydrophilic GGKKKKK sequence on signaling has also been determined by generating a solubility tag peptide (LGGKKKKK). 4. The main figures have been modified to include the overall structure of PC1 protein.

