## Supplementary material for "Activation of Polycystin-1 Signaling by Binding of Stalk-derived Peptide Agonists": Table S1, Table S2, Table S3, Table S4, Table S5, Fig. S1, Fig. S2, Fig. S3, Fig. S4, Fig. S5, Fig. S6, Fig. S7, Fig. S8, Fig. S9, Fig. S10

[1]Center for Computational Biology and Department of Molecular Biosciences, University of Kansas, Lawrence, KS 66047; [2]Departments of Biochemistry and Molecular Biology, [3]Clinical Laboratory Sciences, [4]The Jared Grantham Kidney Institute, University of Kansas Medical Center, Kansas City, KS 66160, [5]Dept of Physics, and Center for Biophysics and Computational Biology, Temple University, Philadelphia, PA 19122, , and [6]Department of Pharmacology and Computational Medicine Program, University of North Carolina – Chapel Hill, Chapel Hill, NC 27599.

* Corresponding

### Experimental procedures

#### Antibodies and peptide synthesis

Primary antibodies used included A19, for detection of mouse PC1 CTF [1], and TRAM2 (Epitomics, 3685-1). Secondary antibodies conjugated to HRP were purchased from Sigma or Jackson ImmunoResearch. Stalk-derived peptides were synthesized by GenScript using the Fmoc method and verified by HPLC-MS analysis.

#### DNA expression constructs and cloning

To produce CTF expression construct of mouse (m) PC1, sequences starting at T3041 and proceeding past the first TM domain were amplified by PCR from PC1-11TM [2], respectively, using 5'-mCleavStalkBsm-For and 3'-TMI-EcoRV primers to produce mCleavStk, which was joined via the BsmBI site to a PCR product encoding the signal peptide sequence (MPMGSLQPLATLYLLGMLVASVLG) from the T cell surface glycoprotein CD5[3] in pBlueScript (pBS) to generate pBS-mCD5-cleaved stalk. The EcoRI-EagI fragment containing mCD5-cleaved stalk was joined to a 3.2 kb EagI-NotI fragment from PC1-11TM encoding TM2 through the C-tail to produce the final pCIneo-mCTF expression construct. The stalkless CTF mutant expression construct, pCIneo-mCTF$^{\Delta st}$, starting with S3062 of the mouse PC1 stalk was generated by the same scheme except for using the 5'-mΔStalkBsmFor primer for the initial PCR. PCR and mutagenesis primers were synthesized by IDT and sequences are listed in **Table S5**. PCR and mutagenesis products and their final constructs were confirmed by Sanger sequencing (GeneWiz). Expression constructs of CTF and CTF$^{\Delta st}$ from human PC1 were made in pCI vector as described previously[10]. In the conduct of research utilizing recombinant DNA, the investigator adhered to NIH Guidelines for research involving recombinant DNA molecules.

### Cell culture and transient transfection

HEK293T cells (ATCC) were maintained and transiently transfected as described previously [4]. Cells were passaged into 6-well plates (6 X $10^5$ cells/well; 3 wells/transfection condition) and transfected with a DNA mixture containing either the 4xNFAT or the 7xAP-1 promoter-Firefly luciferase reporter (100 ng; Stratagene), along with Renilla luciferase (50 ng of pGL4.70[*hRluc*] or 1 ng of pRL-null; Promega), and pCI expression vector encoding either CTF or $CTF^{\Delta st}$ (75 ng for signaling; 600 ng for surface biotinylation) or an equimolar amount of empty pCI vector as control. pBlueScript (Stratagene) was used to bring the total DNA amount to 8 ug. After 2.5 hrs, the DNA mixture was replaced with serum-free culture medium, and after 20-24 hrs, cells were lysed in 1X Passive Lysis buffer (PLB; Promega) supplemented with protease and phosphatase inhibitors. Firefly (Fluc)- and Renilla (Rluc)-luciferase activity in each cell lysate was determined using the Dual Luciferase Assay Kit (Promega) and a Berthold tube luminometer. NFAT-Fluc luminescence was normalized to Rluc for each well within a transfection condition, and then averaged for each condition (n =3 wells/condition). Means of normalized NFAT-Fluc with standard deviation [5] were graphed. Signaling-transfection experiments were performed a minimum of 3 times (i.e., $\geq$ 3 biological replicates) each with 3 technical replicates/condition except where noted.

### Stalk peptide treatment

Cells were plated into 24-well plates (1.5 X $10^5$ cells/well) and transfected with $CTF^{\Delta st}$ or empty pCI expression vectors, along with NFAT-Fluc and Rluc plasmids. Two hours following medium exchange, one-half of culture medium volume was replaced with an equal volume of either serum-free medium (no peptide control), or stalk-derived or solubility tag peptide (2 mM in serum-free medium) and incubated overnight. In some experiments, an additional 50-100 ul of peptide (1 mM) was added the following morning. Cells were lysed at 24 hrs following the initial peptide or control medium addition.

**Cell surface biotinylation analyses**

Surface labeling [6] was performed on intact cells 22-24 hrs post-transfection using 1.5 mg/ml PBS of the membrane-impermeable, cleavable biotin cross-linking reagent (Sulfo-NHS-SS-Biotin; Pierce) for 30 min on ice. Crosslinking was inactivated by addition of 50 mM Tris, pH 8.0. Cells were washed and then lysed in 1X PLB with protease inhibitors. A 10% aliquot of the total cell lysate was removed and saved as the input sample. NeutraAvidin-agarose beads (Pierce) were added to remove biotinylated surface proteins. The supernatant was removed and saved as the unbound cytosolic fraction (sup). A representative amount of each fraction, i.e., the total lysate (input), cytosolic (sup), and biotinylated surface proteins (beads) was analyzed by SDS-PAGE/Western blot. TRAM2 (ER resident protein) was used as a cell fractionation marker.

**Western blot analyses**

Gel loading volumes of signaling cell lysates were calculated based on normalization to relative Rluc activity within a condition.  Lysate samples were electrophoresed through 7.5% denaturing polyacrylamide gels and transferred to nitrocellulose membrane using the TransBlot Turbo and ReadyBlot transfer buffer (BioRad). Blots were incubated with primary antibody (1:1,000 dilution for A19; 1:10,000 for anti-TRAM2) in Tris-buffered saline (TBS; 10 mM Tris pH 7.4, 150 mM NaCl) with 0.1% Tween-20 (TBST) and 5% powdered dry milk for 14-16 hrs at 4˚C. Secondary antibodies conjugated to HRP were incubated at 1:5,000 dilution in TBST/5% milk for 1 hr at room temperature. Blots were developed using a chemiluminescent substrate (Clarity; BioRad), and multiple exposures were captured for each blot with the Amersham Imager 600 with the band saturation detection mode enabled. Volume density (minus background) of immunoreactive bands was determined using ImageQuant TL (GE Healthcare). In most cases, duplicate blots were prepared and quantified to obtain an average band density relative to wild type CTF.

#### Statistical analyses

Statistical analysis was performed using GraphPad Prism 9 (GraphPad Software, San Diego, CA, USA). Data are presented as means + standard deviation [5] for bar graphs. Multiple comparisons used one-way ANOVA and Tukey's post-hoc analysis. $P \leq 0.05$ was considered statistically significant.

### Computational Methods

#### Gaussian Accelerated Molecular Dynamics (GaMD)

GaMD is an unconstrained enhanced sampling approach that works by adding a harmonic boost potential to smooth the potential energy surface of biomolecules to reduce energy barriers[7]. Brief description of the method is provided here.

Consider a system with $N$ atoms at positions $\vec{r} = \{\vec{r}_1, \ldots, \vec{r}_N\}$. When potential energy of the system $V(\vec{r})$ is less than a threshold energy $E$, a boost potential $\Delta V(\vec{r})$ is added to the system as follows:

$$V^*(\vec{r}) = V(\vec{r}) + \Delta V(\vec{r}), V(\vec{r}) < E \quad (1)$$

$$\Delta V(\vec{r}) = \frac{1}{2}k(E - V(\vec{r}))^2, V(\vec{r}) < E, \quad (2)$$

where $k$ is the harmonic force constant. The two adjustable parameters $E$ and $k$ can be determined by application of three enhanced sampling principles. First, for any two arbitrary potential values $V_1(\vec{r})$ and $V_2(\vec{r})$ found on the original energy surface, if $V_1(\vec{r}) < V_2(\vec{r})$, $\Delta V$ should be a monotonic function that does not change the relative order of the biased potential values, i.e., $V_1^*(\vec{r}) < V_2^*(\vec{r})$. Second, if $V_1(\vec{r}) < V_2(\vec{r})$, the potential difference observed on the smoothed energy surface should be smaller than that of the original, i.e., $V_2^*(\vec{r}) - V_1^*(\vec{r}) < V_2(\vec{r}) - V_1(\vec{r})$. By combining the first two criteria and plugging in the formula of $V^*(\vec{r})$ and $\Delta V$, we obtain:

$$V_{max} \le E \le V_{min} + \frac{1}{k}, \quad (3)$$

where $V_{min}$ and $V_{max}$ are the system minimum and maximum potential energies. To ensure that eq (3) is valid, $k$ has to satisfy: $k \le \frac{1}{V_{max}-V_{min}}$. Let us define $k \equiv \frac{k_0}{V_{max}-V_{min}}$, then $0 < k_0 \le 1$. Third, the standard deviation (SD) of $\Delta V$ needs to be small enough (i.e., narrow distribution) to ensure accurate reweighting using cumulant expansion to the second order: $\sigma_{\Delta V} = k(E - V_{avg})\sigma_V \le \sigma_0$, where $V_{avg}$ and $\sigma_V$ are the average and SD of ΔV with $\sigma_0$ as a user-specified upper limit (e.g., $10k_BT$) for accurate reweighting. When $E$ is set to the lower bound $E = V_{max}$ according to eq 3, $k_0$can be calculated as:

$$k_0 = \min(1.0, k_0') = min\left(1.0, \frac{\sigma_0}{\sigma_V} . \frac{V_{max}-V_{min}}{V_{max}-V_{avg}}\right) \quad (4)$$

Alternatively, when the threshold energy $E$ is set to its upper bound $E = V_{min} + \frac{1}{k}$, $k_0$ is set to:

$$k_0 = k_0^{"} \equiv \left(1 - \frac{\sigma_0}{\sigma_V}\right) . \frac{V_{max}-V_{min}}{V_{avg}-V_{min}} \quad (5)$$

if $k_0^{"}$ is calculated between 0 and 1. Otherwise, $k_0$ is calculated using Eq. (4).

**Peptide Gaussian accelerated molecular dynamics (Pep-GaMD)**

Peptides often undergo large conformational changes during binding to target proteins, being distinct from small-molecule ligand binding or protein-protein interactions (PPIs). In this regard, Peptide GaMD or "Pep-GaMD" has been developed to enhance sampling of peptide binding[9]. In Pep-GaMD, we consider a system of peptide *L* binding to a protein *P* in a biological environment *E*. Presumably, peptide binding mainly involves in both the bonded and non-bonded interaction energies of the peptide since peptides often undergo large conformational changes during binding to the target proteins. Thus, the essential peptide potential energy is $V_L(r) = V_{LL,b}(r_L) + V_{LL,nb}(r_L) + V_{PL,nb}(r_{PL}) + V_{LE,nb}(r_{LE})$. In Pep-GaMD, we add boost potential selectively to the essential peptide potential energy according to the GaMD algorithm:

$$\Delta V_L(r) = \begin{cases} \frac{1}{2} k_L \left(E_L - V_L(r)\right)^2, & V_L(r) < E_L \\ 0, & V_L(r) \geq E_L \end{cases} \quad (6)$$

where $E_L$ is the threshold energy for applying boost potential and $k_L$ is the harmonic constant. In addition to selectively boosting the peptide, another boost potential is applied on the protein and solvent to enhance conformational sampling of the protein and facilitate peptide rebinding. This boost represents the total system potential energy without the essential peptide potential energy included:

$$\Delta V_D(r) = \begin{cases} \frac{1}{2} k_D \left(E_D - V_D(r)\right)^2, & V_D(r) < E_D \\ 0, & V_D(r) \geq E_D \end{cases} \quad (7)$$

Where $V_D$ represents the total system potential energy without the essential peptide potential energy included, $E_D$ represents the second boost potential threshold energy and $k_D$ represents the harmonic constant. Hence, this contributes to the dual-boost Pep-GaMD as the total boost potential $\Delta V(r) = \Delta V_L(r) + \Delta V_D(r)$.

**Energetic Reweighting of Pep-GaMD Simulations**

For energetic reweighting of Pep-GaMD simulations to calculate potential mean force (PMF), the probability distribution along a reaction coordinate is written as $p^*(A)$. Given the boost potential $\Delta V(r)$ of each frame, $p^*(A)$ can be reweighted to recover the canonical ensemble distribution $p(A)$, as:

$$p(A_j) = p^*(A_j)\frac{\langle e^{\beta \Delta V(r)}\rangle_j}{\sum_{i=1}^{M}\langle p^*(A_i) e^{\beta \Delta V(r)}\rangle_i}, \quad j = 1,\dots, M \tag{8}$$

where $M$ is the number of bins , $\beta = k_B T$ and $\langle e^{\beta \Delta V(r)}\rangle_j$ is the ensemble-averaged Boltzmann factor of $\Delta V(r)$ for simulation frames found in the $j^{\text{th}}$ bin. The ensemble-averaged reweighting factor can be approximated using cumulant expansion:

$$\langle e^{\beta \Delta V(r)}\rangle_j = \exp\left\{\sum_{k=1}^{\infty}\frac{\beta^k}{k!}C_k\right\} \tag{9}$$

where first two cumulants are given by

$$C_1 = \langle \Delta V \rangle$$

$$C_2 = \langle \Delta V^2 \rangle - \langle \Delta V \rangle^2 = \sigma_V^2 \tag{10}$$

The boost potential obtained from Pep-GaMD simulations usually follows near-Gaussian distribution. Cumulant expansion to the second order thus provides a good approximation for computing the reweighting factor. The reweighted free energy $F(A) = -k_B T \ln p(A)$ is calculated as

$$F(A) = F^*(A) - \sum_{k=1}^{2}\frac{\beta^k}{k!}C_k + F_c \tag{11}$$

where $F^*(A) = -k_B T \ln p^*(A)$ is the modified free energy obtained from GaMD simulation and $F_c$ is a constant.

### Computational model of peptide agonist-bound ΔStalk PC1 CTF and HPEPDOCK docking

With GaMD simulations of the WT PC1 CTF obtained from the previous study that revealed a active TA/stalk-mediated allosteric signaling[10], structural clustering of the extracellular regions of PC1 CTF, including the Stalk, TM2-TM3 and TM4-TM5 loops, TOP domain, S3-S4 loop and pore loop, was performed using the hierarchical agglomerative algorithm in CPPTRAJ[11]. The top-ranked representative conformation of PC1 CTF was used for peptide docking after removal of the TA/stalk (ΔStalk). Then the HPEPDOCK[12] webserver was applied to dock the p9, p17 and p21 stalk-derived peptides to ΔStalk CTF.

### Simulation system setup

We embedded ΔStalk CTF in a palmitoyl-oleoyl-phosphatidyl-choline (POPC) bilayer and solvated the system in 0.15 M NaCl explicit solvent using CHARMM-GUI (**Fig. S3C**). Neutral patches (acetyl and methylamide) were added to the protein termini residues. The peptide termini were kept as charged (NH3+ and COO-). The CHARMM36m[13] force field parameters were used for the protein, peptides and lipids. CHARMM-GUI output files and scripts were used with default parameters to prepare the systems for Pep-GaMD simulations. Energy minimization was performed for 5000 steps using constant number, volume and temperature (NVT) ensemble at 310 K. Further equilibration was done for 375 ps at 310 K using NPT ensemble. Conventional MD (cMD) simulations was performed on the systems for 10 ns at 1 atm pressure and 310 K temperature. All-atom Pep-GaMD simulations were performed with a short cMD for 10 ns, Pep-GaMD equilibration for 55 ns followed by three independent Pep-GaMD production runs for 500 ns for each system with randomized initial atomic velocities. A cutoff distance of 9 Å was used for the van der Waals and short-range electrostatic interactions, and long-range electrostatic interactions were computed with the particle-mesh Ewald summation method [14]. The simulation systems are ~90 × 136 × 117Å$^3$ in dimension, containing a total of ~100 K atoms with explicit solvent and lipid molecules.

**Simulation Analysis**

Pep-GaMD simulation trajectories were analyzed using CPPTRAJ [15] and VMD [16] tools. Trajectory analysis showed peptides binding the TOP domain of PC1 CTF. A previously identified salt bridge interaction between the TOP domain R3848 and the pore loop E4078 of the PC1 protein, that is an important interaction during PC1 signal activation. The number of contacts formed between peptides p9, p17 and p21, respectively, and the R3848-E4078 salt bridge distance were used as reaction coordinates to calculate 2D free energy profiles using the *PyReweighting* toolkit [17]. A bin size of 2 Å was used for distances and 10 for the number of contacts. Three independent Pep-GaMD simulations were combined to perform structural clustering using the hierarchical agglomerative clustering algorithm in CPPTRAJ[15]. A 3 Å RMSD cutoff was used for each peptide system. PyReweighting[17] was then applied to calculate the original free energy values of each peptide structural cluster with a cutoff of 500 frames. The structural clusters were finally ranked according to the reweighted free energy values.

### Potts sequence covariation model

Using a seed alignment of 189 orthologs of human PKD1 obtained from the Ensemble database [18], we used iterative searching of the Uniprot Database using HHblits[19] to obtain a multiple-sequence alignment of 4384 homologs, and after filtering using standard methods with an 80% identity threshold [20] we obtained 1022 effective sequences of length 853. These sequences had an average of 23% sequence identity reflecting extensive diversity across Eukaryotes. As this number of sequences and sequence length could lead to an overfit model[21], we used a subset the MSA to limited to 394 positions on the extracellular side of PKD1 including the GAIN domain, stalk, TOP domain, and ends of the transmembrane helices. We inferred a Potts model from this reduced MSA using the Mi3-GPU software [20]. We evaluated the position-pair statistical interactions using the "weighted frobenius norm" interaction score[20], and computed the residue-residue covariation values $C_{ab}^{ij} = f_{ab}^{ij} - f_a^i f_b^j$ as the difference between the pair-residue frequency $f_{ab}^{ij}$ of letters a,b at positions i,j and the null expectation under assumption of site-independence by multiplying the two single-site frequencies, $f_a^i$ and $f_b^j$. The maximum possible covariance is 25%. Statistically significant covariations scores will be greater than the expected binomial sampling error given our dataset size of N~1022, and for bivariate count $f_{ab}^{ij}$ the binomial-sampling standard deviation is $\sqrt{f_{ab}^{ij}(1 - f_{ab}^{ij})/N}$ which for the typical bivariate frequency of ~10% corresponds to a ~1% error. To choose a cutoff in Frobenius Norm to distinguish likely contacts from noise, we compared the Potts interactions scores to the contacts predicted using Alphafold[22] structure, choosing the plotting cutoff at false-positive rate of 50% relative to the contacts predicted using the Alphafold structure using an 8Å nearest-heavy atom side chain distance.

**Peptide binding free energy calculations**

Molecular Mechanics/Poisson-Boltzmann Surface Area (MM/PBSA) analysis was performed to calculate the binding free energies of peptides p9, p17 and p21 to PC1 CTF. The analysis was performed using the trajectory in which the peptide was bound to the receptor. In MM/PBSA[24], the binding free energy of the ligand (L) to the receptor (R) to form the complex (RL) is calculated as:

$$\Delta G_{bind} = G_{RL} - G_R - G_L \tag{12}$$

where $G_{RL}$ is the Gibbs free energy of the complex RL, $G_R$ is the Gibbs free energy of the molecule R in its unbound state and $G_L$ is the Gibbs free energy of the molecule L in its unbound state, respectively.

$\Delta G_{bind}$ can be divided into contributions of different interactions as[25]:

$$\Delta G_{bind} = \Delta H - T\Delta S = \Delta E_{MM} + \Delta G_{sol} - T\Delta S \tag{13}$$

in which

$$\Delta E_{MM} = \Delta E_{int} + \Delta E_{elec} + \Delta E_{vdW} \tag{14}$$

$$\Delta G_{sol} = \Delta G_{PB/GB} + \Delta G_{SA} \tag{15}$$

$$\Delta G_{SA} = \gamma . SASA + b \tag{16}$$

where $\Delta E_{MM}$ , $\Delta G_{sol}$ , $\Delta H$ and $-T\Delta S$ are the changes in the gas-phase molecular mechanics (MM) energy, solvation free energy, enthalpy and conformational entropy upon ligand binding, respectively. $\Delta E_{MM}$ includes the changes in the internal energies $\Delta E_{int}$ (bond, angle and dihedral energies), electrostatic energies $\Delta E_{elec}$ , and the van der Waals energies $\Delta E_{vdW}$. $\Delta G_{sol}$ is the sum of the electrostatic solvation energy $\Delta G_{PB/GB}$ (polar contribution) and the nonpolar contribution $\Delta G_{SA}$ between the solute and the continuum solvent. The polar contribution is calculated using either the Poisson Boltzmann (PB) or Generalized Born (GB) model, while the nonpolar energy is usually estimated using the solvent-accessible surface area (SASA)[26,27] where $\gamma$ is surface tension coefficient and b is the constant offset. The change in conformational entropy $-T\Delta S$ is usually calculated by normal-mode analysis[25] on a set of conformational snapshots taken from MD simulations. However, due to the large computational cost, changes in the conformational entropy are usually neglected as we were concerned more on relative binding free energies of the similar peptide ligands.

MM/PBSA analysis was performed using the gmx_MMPBSA[28] software with the following command line:

```
gmx_MMPBSA -O -i mmpbsa.in -cs com.tpr -ci index.ndx -cg 1 13 -ct com_traj.xtc -cp topol.top -o FINAL_RESULTS_MMPBSA.dat -eo FINAL_RESULTS_MMPBSA.csv
```

Input file for running MM/PBSA analysis:

```
&general
sys_name="Prot-Pep-CHARMM",
startframe=1,
endframe=200,
# In gmx_MMPBSA v1.5.0 we have added a new PB radii set named charmm_radii.
# This radii set should be used only with systems prepared with CHARMM force fields.
# Uncomment the line below to use charmm_radii set
#PBRadii=7,
/
&pb
# radiopt=0 is recommended which means using radii from the prmtop file for both the PB calculation and for the NP
# calculation
istrng=0.15, fillratio=4.0, radiopt=0
```

Residue-wise interaction energy analysis was performed on peptides p9, p17 and p21 using the trajectory in which the peptide was bound to the PC1 CTF using the gmx_MMPBSA[28] software with the following command line:

```
gmx_MMPBSA -O -i mmpbsa.in -cs com.tpr -ct com_traj.xtc -ci index.ndx -cg 3 4 -cp topol.top -o FINAL_RESULTS_MMPBSA.dat -eo FINAL_RESULTS_MMPBSA.csv -do FINAL_DECOMP_MMPBSA.dat -deo FINAL_DECOMP_MMPBSA.csv
```

Input file for running residue-wise energy decomposition analysis:

```
&general
sys_name="Decomposition",
startframe=1,
endframe=200,
#forcefields="leaprc.protein.ff14SB"
/
&gb
igb=5, saltcon=0.150,
/
#make sure to include at least one residue from both the receptor
#and peptide in the print_res mask of the &decomp section.
#this requirement is automatically fulfilled when using the within keyword.
#http://archive.ambermd.org/201308/0075.html
&decomp
idecomp=2, dec_verbose=3,
print_res="A/854-862 A/1-853”,
/
```

**Table S1. Summary of MM/PBSA binding free energy analysis for the peptides p9, p17 and p21 and PC1 CTF in the bound state sampled during Pep-GaMD simulations.**

| System | *Δ*G (kcal/mol) |
|---|---|
| p21 | -40.29±6.94 |
| p9 | -17.30±4.50 |
| p17 | -12.74±5.62 |

**Table S2. Summary of residue-wise energy decomposition analysis between the peptide p9 and PC1 CTF in the bound state sampled during Pep-GaMD simulations.**

| Residue | *Δ*G (kcal/mol) |
|---|---|
| **T1** | **-10.47±6.92** |
| A2 | -3.28±2.18 |
| F3 | -2.62±2.62 |
| G4 | -0.30±2.38 |
| A5 | -1.61±2.67 |
| S6 | -0.59±4.49 |
| L7 | -0.28±2.96 |
| F8 | -1.51±1.84 |
| **V9** | **-10.14±5.68** |

| | |
|---|---|
| R3891 | -0.14±7.26 |
| R3892 | -0.31±7.85 |
| **F3888** | **-3.24±2.95** |
| H3864 | -0.03±3.86 |
| R3970 | -0.94±2.62 |
| R3968 | -0.27±1.67 |

**Table S3. Summary of residue-wise energy decomposition analysis between the peptide p17 and PC1 CTF in the bound state sampled during Pep-GaMD simulations.**

| **Residue** | **ΔG (kcal/mol)** |
|---|---|
| **T1** | **-10.98±2.32** |
| A2 | -0.36±2.34 |
| F3 | -0.23±3.82 |
| G4 | -0.03±3.83 |
| A5 | -0.86±3.08 |
| S6 | -0.48±4.06 |
| L7 | -0.05±2.90 |
| F8 | -0.21±4.42 |
| V9 | -0.19±3.41 |
| P10 | -0.72±2.64 |
| P11 | -0.09±3.23 |
| S12 | -0.29±4.92 |
| H13 | -1.19±7.37 |
| V14 | -1.25±3.08 |
| **R15** | **-10.63±3.76** |
| F16 | -0.93±5.52 |
| **V17** | **-10.20±3.83** |
| Y3307 | -0.11±2.68 |
| H3311 | -0.04±3.25 |
| **R3314** | **-8.90±1.65** |
| **R3700** | **-14.63±7.8** |
| Q3707 | -0.27±4.96 |
| S3711 | -0.08±4.06 |
| **E3708** | **-10.14±4.15** |

**Table S4. Summary of residue-wise energy decomposition analysis between the peptide p21 and PC1 CTF in the bound state sampled during Pep-GaMD simulations.**

| **Residue** | **ΔG (kcal/mol)** |
|---|---|
| T1 | -0.25±1.66 |
| A2 | -1.15±1.31 |
| F3 | -1.02±2.90 |
| G4 | -0.08±0.10 |
| A5 | -2.42±2.65 |
| S6 | -0.11±2.15 |
| L7 | -0.09±1.31 |
| F8 | -0.83±1.43 |
| V9 | -0.68±1.10 |
| **P10** | **-8.17±2.10** |
| **P11** | **-6.16±1.79** |
| S12 | -1.22±2.83 |
| H13 | -0.05±2.56 |
| V14 | -1.13±3.74 |
| R15 | -1.16±1.78 |
| F16 | -0.59±1.02 |
| V17 | -1.59±1.13 |
| F18 | -0.20±1.37 |
| **P19** | **-7.09±1.51** |
| E20 | -3.76±2.24 |
| **P21** | **-8.43±2.01** |
| L3863 | -0.33±2.28 |
| L3701 | -0.04±2.39 |
| I3705 | -0.51±2.16 |
| **E3708** | **-1.68±3.11** |
| **R3712** | **-3.85±1.56** |
| F3714 | -0.01±2.28 |
| **W3726** | **-2.93±2.11** |
| H3729 | -0.56±2.00 |
| L3732 | -0.26±2.49 |
| P3733 | -0.61±2.33 |
| N3738 | -0.07±4.91 |
| S3741 | -0.03±4.27 |

**Table S5. Primer Sequences for PCR Cloning and Mutagenesis**.

| Primer | Sequence |
|---|---|
| 5’-CD5 Eco | 5’-_TTCTAGAATTCCCTCGACCTCG -3’ |
| 3’-CD5-BsmBI | 5’- GACTAGCGTCTCATGCCTAGCACGGAAGC -3’ |
| mCleavStalkBsm-For | 5’- GACTAGCGTCTCAGGCACTGCCTTCGGTGCC-3’ |
| mΔStalkBsmFor | 5'- GACTAGCGTCTCAGGCAGTGCAAGCATCAACTACATTGTCC -3' |
| TMI-EcoRV | 5’-GACTAGGATATCCCTCTGGACTCTAGTAAAGCG-3’ |
| BsrGIstalk-Rev | 5’- AGGGTCTGGGTAGAGTGCTT -3’ |

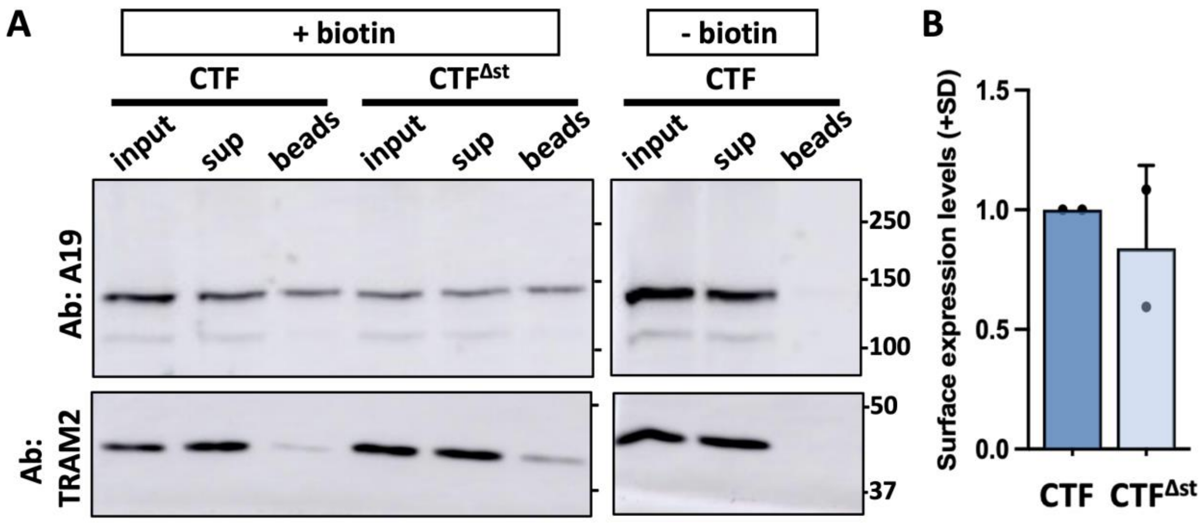


**Figure S1. Cell surface expression of mouse PC1 CTF and mCTF$^{\Delta stk}$.** Representative Western blot of a surface biotinylation experiment with mCTF and mCTF$^{\Delta stk}$. A '- biotin' control was included using CTF-transfected cells for which the NHS-biotin reagent was omitted from the procedure. Aliquots of the total cell lysate (input; after biotinylation and before neutravidin pulldown), supernatant (sup; following neutravidin bead removal), and biotinylated cell surface proteins bound to neutravidin beads (beads) representing 10%, 10% and 70% of each sample, respectively, were analyzed. Blots were probed with A19 and then stripped and reprobed for NaKATPase or the resident ER protein, TRAM2. (**E**) Summary of cell surface expression levels of CTF$^{\Delta stk}$ relative to CTF (means +SD) from two separate experiments.

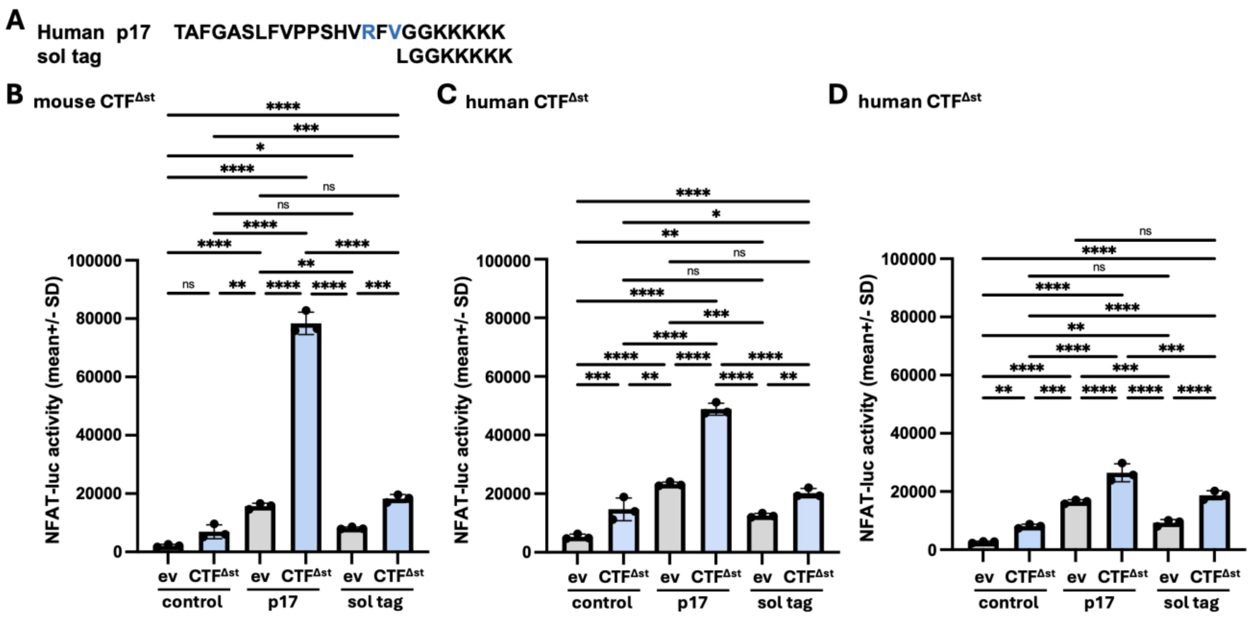


**Figure S2. Solubility tag peptide treatment of ev- or CTF$^{\Delta st}$-transfected cells. (A)** Sequences of the p17 stalk peptide derived from human PC1 and the solubility tag peptide (sol tag). Residues differing from mouse PC1 stalk sequence are shown in blue. **(B-D)** HEK293T cells were transfected with empty expression vector (ev) or the CTF$^{\Delta st}$ expression construct from either mouse (B) or mouse PC1, and transfected cells were treated either without (negative control; culture medium only) or with human p17 peptide (positive control) or sol tag peptide. Graphs show the NFAT-luc activity for ev- (gray bars) and CTF$^{\Delta st}$- (blue bars) transfected cells after 24 hr treatment with or without peptide for each experiment. Results are the means (+SD) of reporter activity from 3 wells/condition in each experiment. *, $p < 0.05$, **, $p < 0.01$, ***, $p < 0.001$; ****, $p < 0.0001$; ns = not significant. Analysis by 1-way ANOVA with Tukey-Kramer post-test. In each of the experimental replicates, treatment with the sol tag peptide led to a nominal ~2-fold increase in reporter activity with CTF$^{\Delta st}$- versus ev-transfected cells (2.24-, 1.60-, and 2.02-fold for B-D, respectively). Although the level of NFAT reporter activity stimulated by the positive control p17 in CTF$^{\Delta st}$-transfected cells varied between the 3 experiments, the effect of the sol tag peptide on CTF$^{\Delta st}$- (and ev-) transfected cells was consistent and differed significantly in comparison to $CTF^{\Delta st}$-transfected cells with p17 treatment in all three experiments (regardless of PC1 origin). Furthermore, the NFAT-luc activity of $CTF^{\Delta st}$+sol tag-treated cells did not differ from ev+p17-treated cells for each experiment, suggesting that the 'background' reporter activation observed with ev+stalk peptide versus ev+no peptide control may be due to the solubility tag. Altogether, these results support the rescue of $CTF^{\Delta st}$ signaling by certain stalk-derived peptides is specific to the stalk sequence itself.

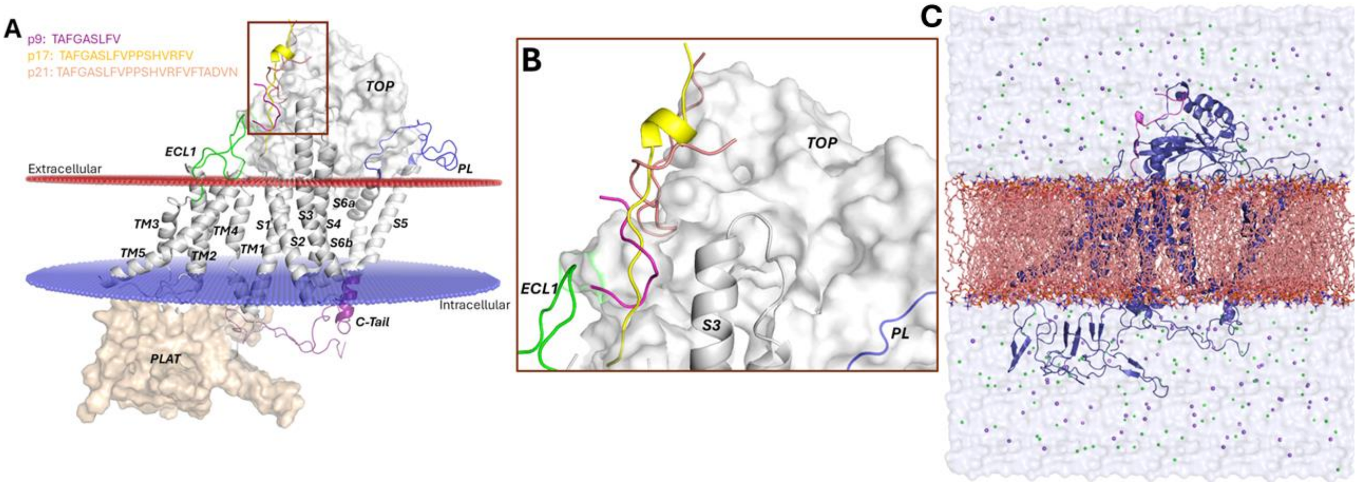


**Figure S3. (A-B**) Docking conformations of p9, p17, and p21 to ΔStalk CTF. (**C**) Pep-GaMD simulation system of ΔStalk PC1 CTF (blue cartoon) embedded in a POPC lipid bilayer (orange sticks) solvated in 0.15 M NaCl solution.

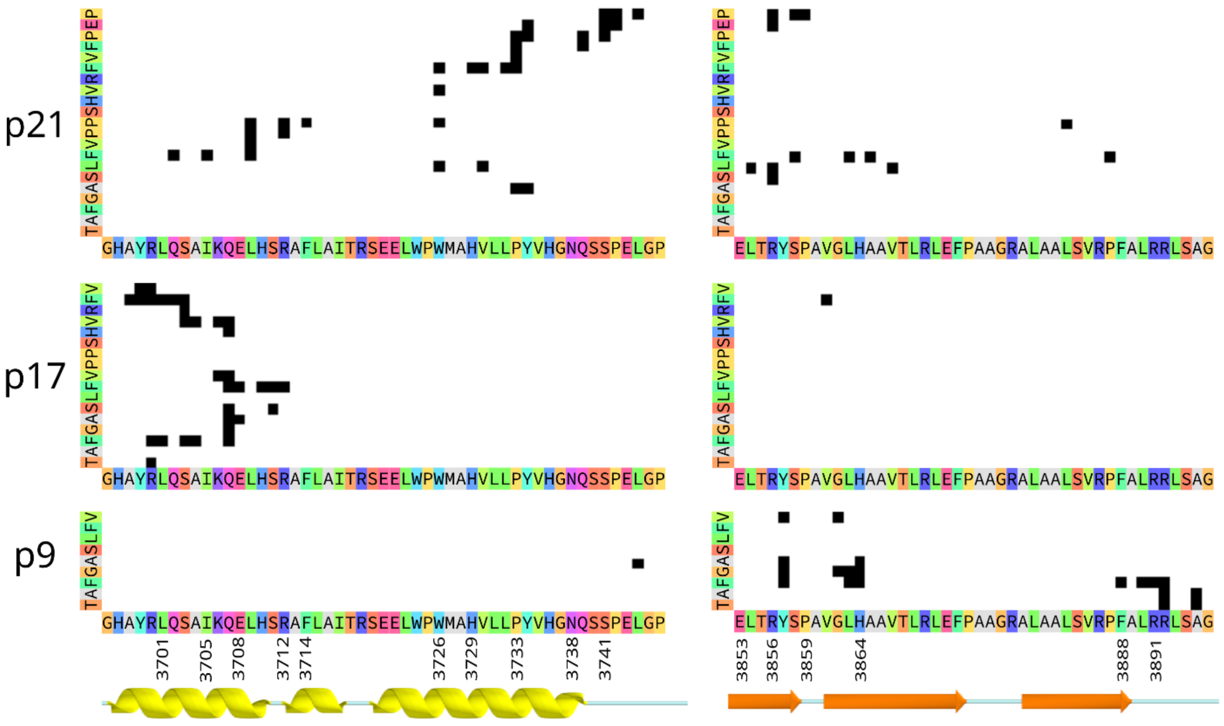


**Figure S4.** Contact maps showing residue-pairs in contact (black squares) between the peptide (y axis) and the extracellular domains of CTF (x-axis), in the representative "Bound" state for p21 (top), p17 (middle) and p9 (bottom). Contacts were defined by a distance less than 4Å between any atom in each residue pair. The secondary structure annotation is colored as in Figure 5, and the sequence is annotated with amino acids in "Taylor" color scheme. For p9, an additional cluster of contacts with residues in helix S3 are not shown.

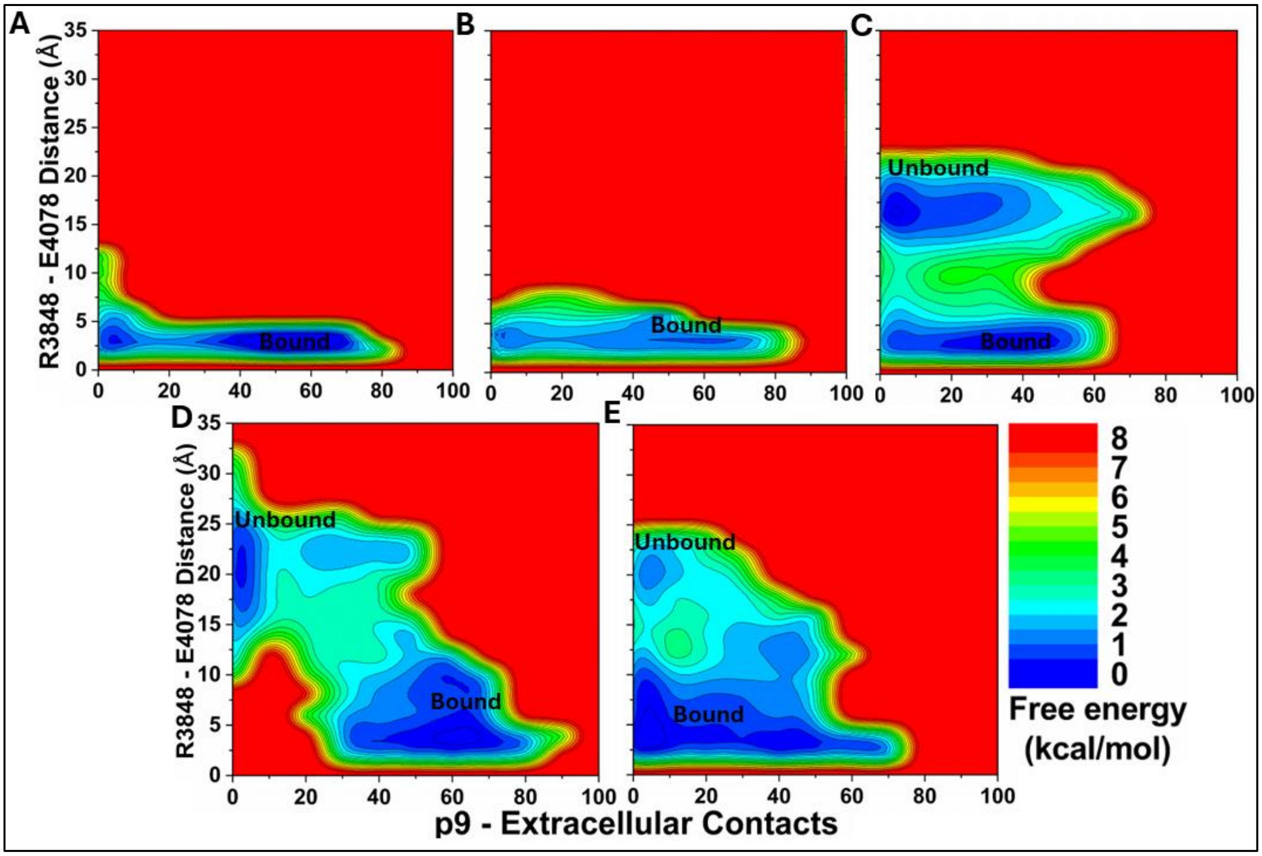


**Figure S5.** 2D free energy profiles of the p9 system regarding the number of atom contacts between the p9 and protein extracellular domains and the R3848-E4078 distance (the CZ atom in R3848 and the CD atom in E4078) calculated from (**A**) Sim1, (**B**) Sim2, (**C**) Sim3, (**D**) Sim4, and (**E**) Sim5 of the Pep-GaMD simulations. Important low-energy conformational states are identified, including the "Unbound" and "Bound".

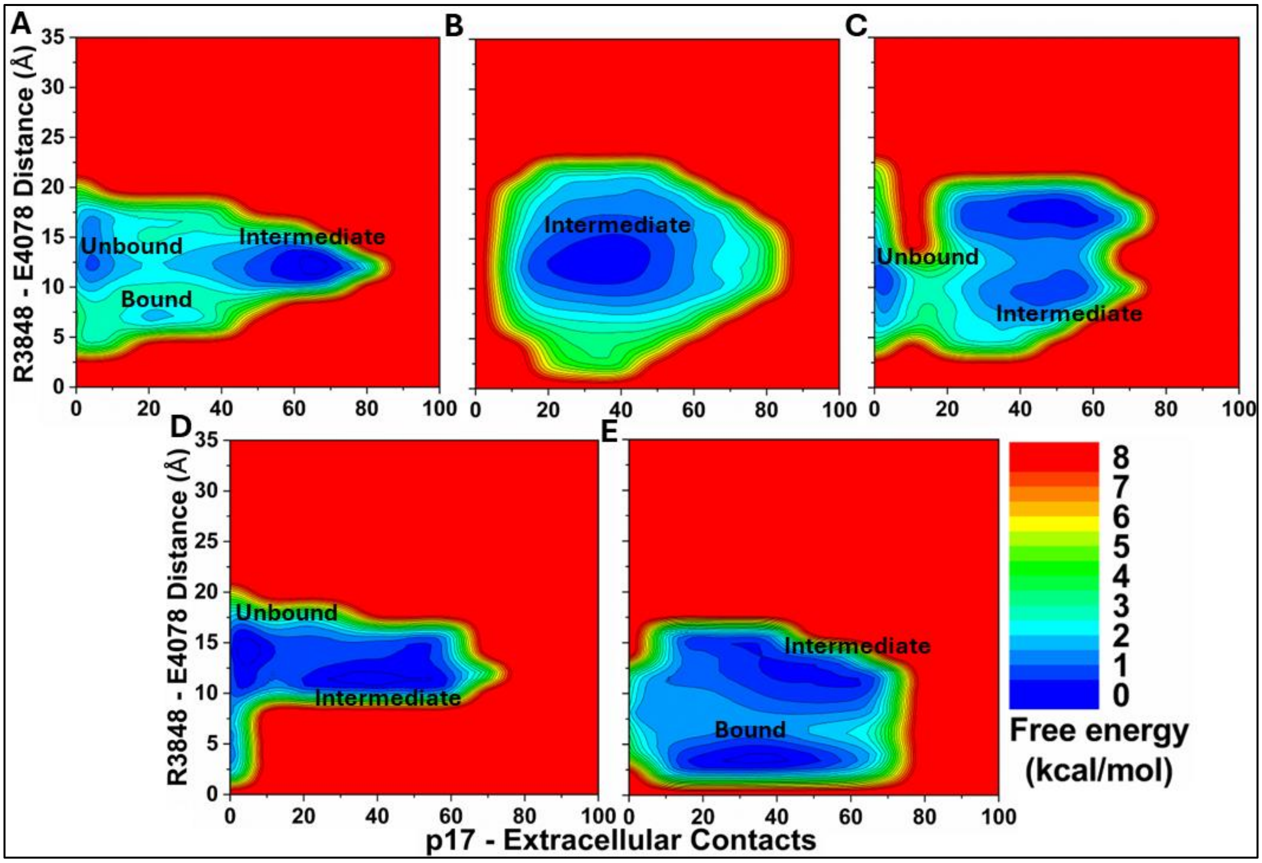


**Figure S6.** 2D free energy profiles of the p17 system regarding the number of atom contacts between the p17 and protein extracellular domains and the R3848-E4078 distance (the CZ atom in R3848 and the CD atom in E4078) calculated from (**A**) Sim1, (**B**) Sim2, (**C**) Sim3, (**D**) Sim4, and (**E**) Sim5 of the Pep-GaMD simulations. Important low-energy conformational states are identified, including the "Unbound" "Intermediate" and "Bound".

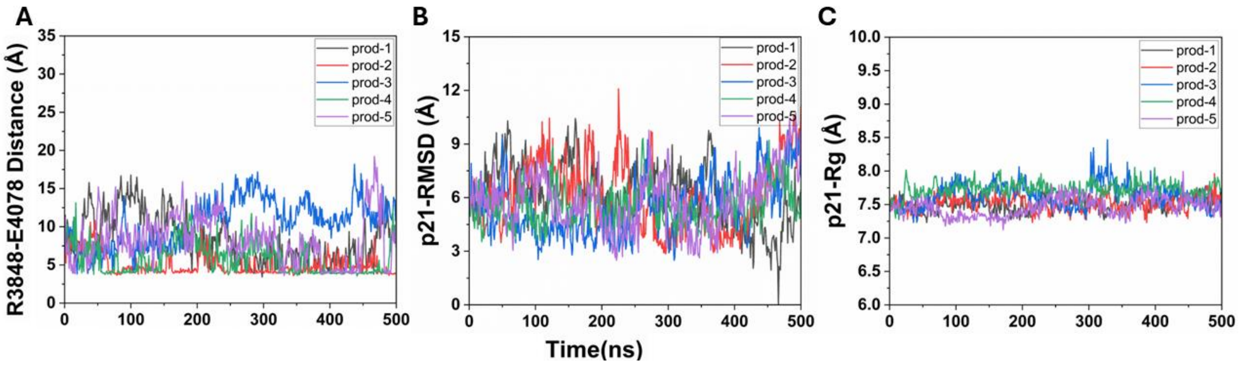


**Figure S7. (A)** Time courses of the p21 system regarding the TOP-PL interaction distance between the CZ atom in R3848 and the CD atom in E4078 calculated from the five Pep-GaMD simulations. **(B)** Time courses of the root-mean square deviation (RMSD) of p21 relative to the starting HPEPDOCK conformation of the peptide calculated from the five Pep-GaMD simulations. **(C)** Time courses of the radius of gyration (*Rg*) of p21 calculated from the five Pep-GaMD simulations.

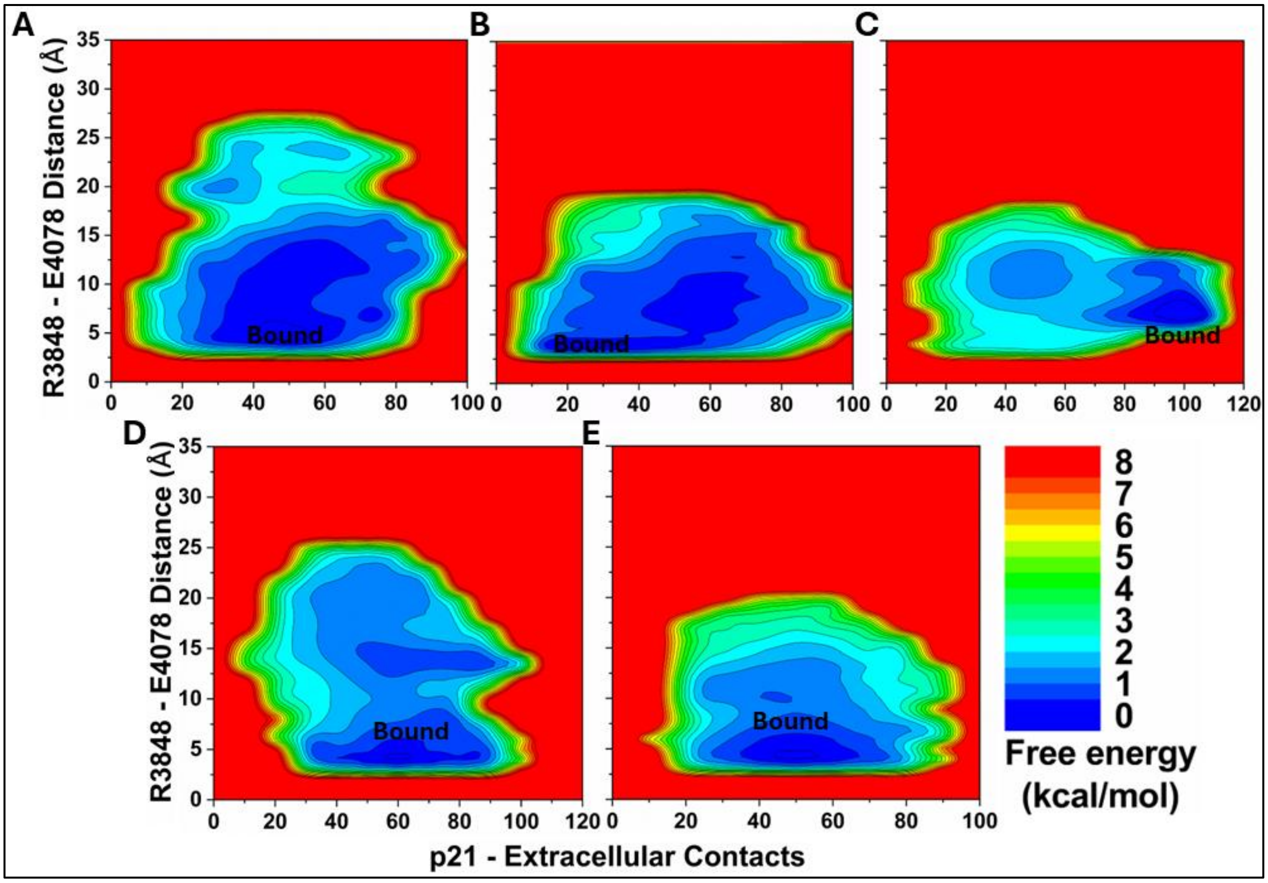


**Figure S8.** 2D free energy profiles of the p21 system regarding the number of atom contacts between the p21 and protein extracellular domains and the R3848-E4078 distance (the CZ atom in R3848 and the CD atom in E4078) calculated from (**A**) Sim1, (**B**) Sim2, (**C**) Sim3, (**D**) Sim4, and (**E**) Sim5 of the Pep-GaMD simulations. Important low-energy conformational state is identified, including the “Bound”.

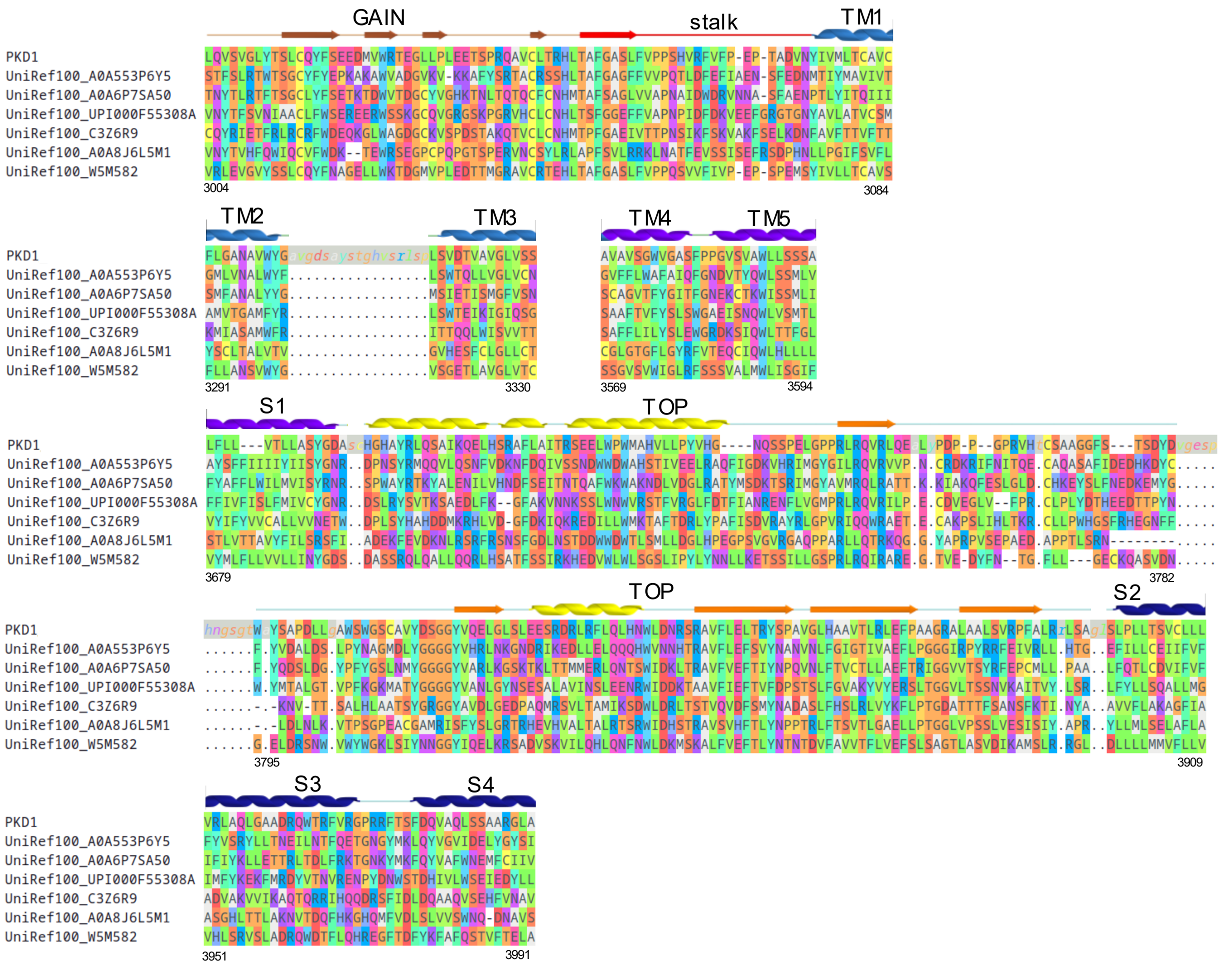


**Figure S9.** Sequence alignment of PKD1 homologs showing the 394-residue extracellular region included in the Potts model. The full alignment includes 4383 sequences, only six diverse homologs are shown for illustration. The PKD1 sequence is shown including inserts relative to the alignment (lower case with gray background, matched with "." in homologs), inserts are not shown for the six homologs. Gaps characters are indicated as "-", representing positions included in the alignment which were missing in that sequence. The human PC1 protein sequence residue numbering is indicated at the bottom ends of each aligned region.

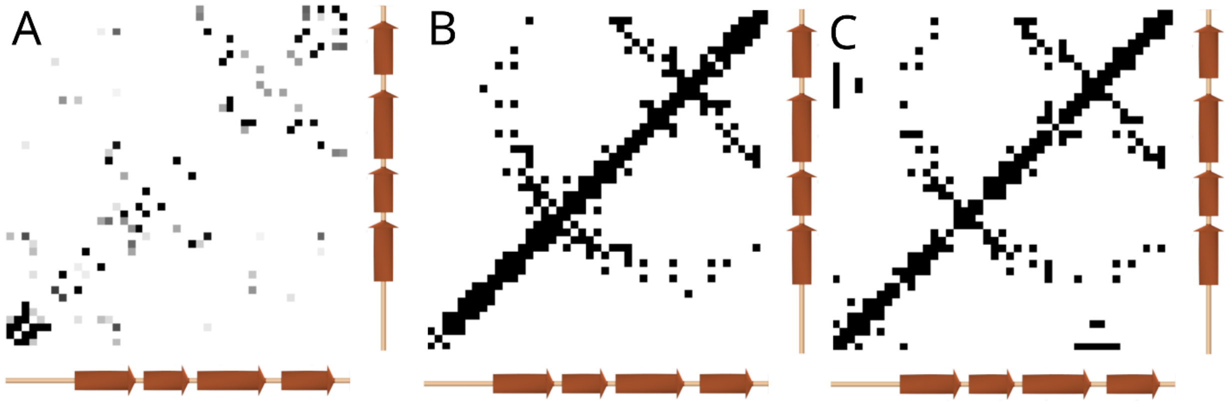


**Figure S10. (A)** Predicted residue-residue interactions within the end of the PC1 GAIN domain using the Potts model, shaded by interaction strength. Secondary structure elements predicted using an Alphafold structure is annotated along the axes. (**B**) Contacts observed in the rat latrophilin-1 GAIN domain crystal structure (PDB: 4DLQ [23]), showing contacts where the nearest side-chain heavy atom distance was under 8Å. (**C**) Contacts observed in the structure of the PKD1 GAIN domain predicted using Alphafold with the same distance criteria. The contacts predicted using the Potts model show correspondence with the structure-based contacts, detecting two antiparallel beta-sheet interactions extending from the diagonal (upper right and middle), plus a possible third antiparallel beta-sheet interface (lower left) not observed in the latrophilin structure as this region may have low similarity.
